# Prefrontal Cortex Astroglia Modulate Anhedonia-like Behavior

**DOI:** 10.1101/2023.05.31.542974

**Authors:** S.A. Codeluppi, M. Xu, Y. Bansal, A.E. Lepack, V. Duric, M. Chow, J. Muir, R.C. Bagot, P. Licznerski, S.L. Wilber, G. Sanacora, E. Sibille, R.S. Duman, C. Pittenger, M. Banasr

## Abstract

Reductions of astroglia expressing glial fibrillary acidic protein (GFAP) are consistently found in the prefrontal cortex (PFC) of patients with depression and in rodent chronic stress models. Here, we examine the consequences of PFC GFAP+ cell depletion and cell activity enhancement on depressive-like behaviors in rodents. Using viral expression of diphtheria toxin receptor in PFC GFAP+ cells, which allows experimental depletion of these cells following diphtheria toxin administration, we demonstrated that PFC GFAP+ cell depletion induced anhedonia-like behavior within 2 days and lasting up to 8 days, but no anxiety-like deficits. Conversely, activating PFC GFAP+ cell activity for 3 weeks using designer receptor exclusively activated by designer drugs (DREADDs) reversed chronic restraint stress-induced anhedonia-like deficits, but not anxiety-like deficits. Our results highlight a critical role of cortical astroglia in the development of anhedonia and further support the idea of targeting astroglia for the treatment of depression.

## INTRODUCTION

Major depressive disorder (MDD) is a severe disorder that has only increased in prevalence in recent years (*1*). Symptoms include feelings of worthless/hopelessness and anhedonia (*2*). MDD is highly comorbid with anxiety disorders (*3*). One of the largest risk factors for MDD is stress and the associated neurophysiological responses resulting from increased allostatic load (*4, 5*). These have drastic effects on the prefrontal cortex (PFC), a region pivotal to the top-down processing of stress (*6*). These effects include reductions in PFC volume, reductions in PFC neuron and astroglial density and size, synaptic loss, and altered dendrite arborization/retraction (*7, 8*). Although differences in the definition of treatment non-response have resulted in highly variable estimates of its prevalence rate. The rates of non-response are commonly reported to be approximately 30% in research settings and up to 55% in the clinic (*9-11*), highlighting the need to investigate the underlying cellular pathologies associated with MDD and to identify new treatment targets.

Astroglia are involved in many brain functions, and their impairment has been implicated in several neurodegenerative and psychiatric pathologies (*12, 13*). Astroglia play an important role in synapse development and function and are a key component of the ‘tripartite synapse’ (*14-16*). Multiple studies have consistently reported astroglial dysfunction in brains from both MDD patients and rodent models of chronic stress, with hallmark findings of decreased astroglial cell density, size, and astroglial specific marker expression in several brain regions, including the PFC (*7, 8, 17-20*). In rodents, a causal relationship between glial loss/deficits was suggested by studies using a gliotoxin to ablate these cells in the PFC; this was sufficient to induce depressive-like behaviors (*21*), increase alcohol preference (*22*), and impair cognitive flexibility (*23*). However, this approach was not selective of astroglia or astroglial subtypes. More specific manipulation is now possible with recent technical advancements in astroglial targeting (*24, 25*).

Among the markers expressed by astroglia, glial fibrillary acidic protein (GFAP) is a key intermediate filament protein involved in maintaining the cytoskeleton structure of a majority of both reactive and non-reactive astroglia in the CNS (*26, 27*). GFAP astroglia are reduced in the PFC of MDD post-mortem (*28*) and rodents after chronic stress (*8*). In contrast, there is no similar reduction in S100b astroglia density, suggesting a specific effect (*29*). Stress-induced GFAP+ cell density changes were associated with GFAP cell atrophy (*20, 30*) and changes in other astroglial proteins (*31, 32*). However, a clear link between the observed reductions in GFAP+ cell density and the development of anxiety- or anhedonia-like behavioral deficits has not been established.

Here, combining genetic and viral approaches, we aimed to establish a causal link between cortical GFAP cell deficits and emotion-related behavior. We investigated the behavioral consequences of selective cortical GFAP+ cell depletion (achieved through artificial expression of the diphtheria-toxin receptor (DTR)) on anxiety, anhedonia, and helplessness. We also tested whether PFC GFAP+ cell activity enhancement (achieved using designer drug exclusively activated by designer drugs [DREADDs]) can reverse the behavioral effects of chronic restraint stress (CRS).

## RESULTS

### Effects of GFAP+ cell depletion on depressive-like behavior

AAV5-GFP-DIOCMV-DTRflag (Fig.1a) aims to induce specific GFAP+ cell depletion. In primary astrocyte cultures, transfected cells expressed green fluorescence protein (GFP) in astrocytes were generated from GFAPcre-pups but DTR fused with flag (DTRflag) in cultures from GFAPcre+ pups (Fig.S1). Following application of diphtheria toxin (DT), we found a 35% reduction of cell survival (*supplementary material*). This construct was then packaged into an AAV5 and used in the follow up studies to examine the consequences of GFAP+ cell depletion on behavior and confirm GFAP+ cell loss (Fig.2).

**Fig. 1:**
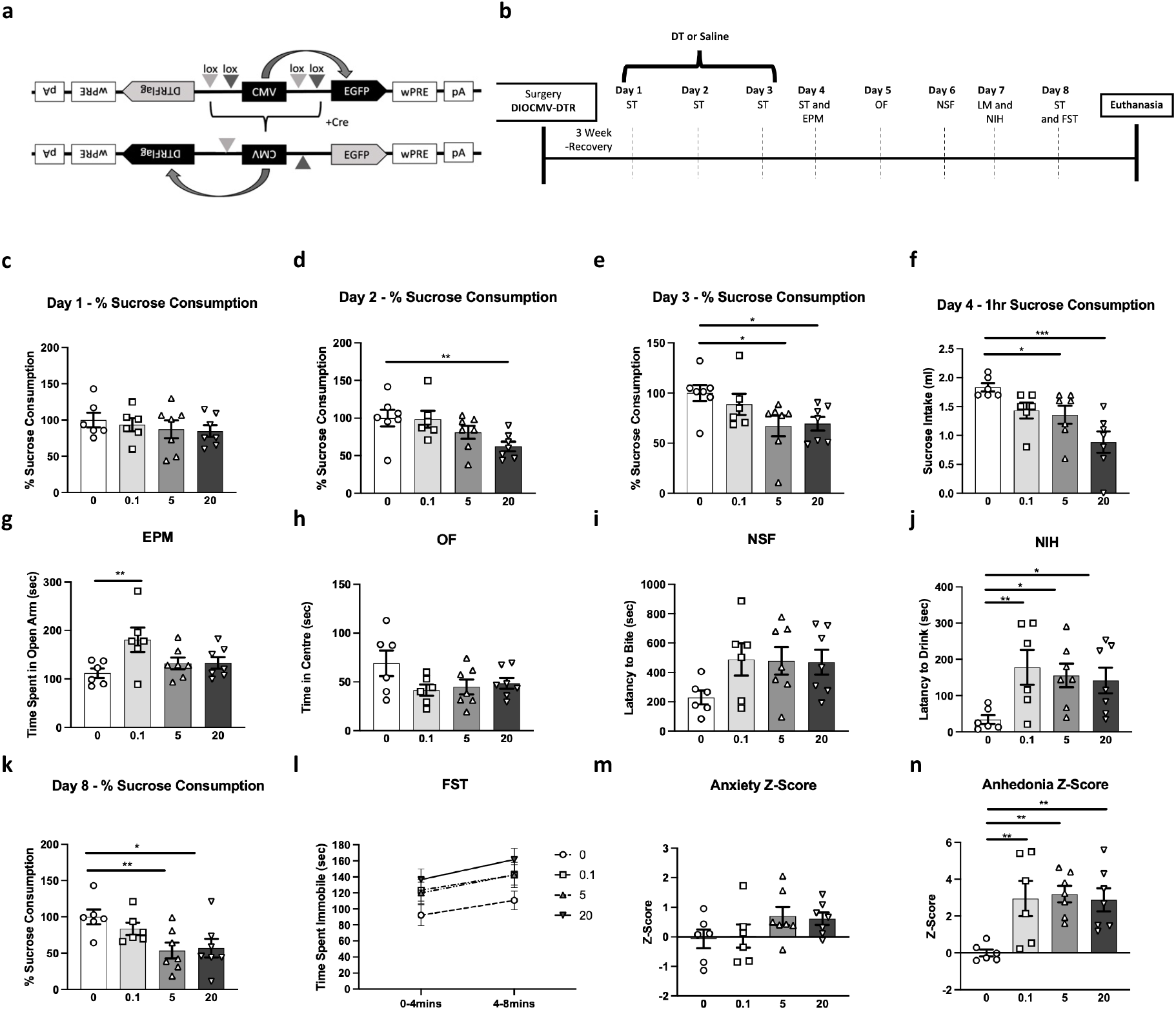
Cortical Glial Fibrillary Acidic Protein (GFAP)+ cell ablation induces behavioral deficits. (a) Schematic representation of the AAV5-GFP-DIOCMV-DTRflag viral construct designed to conditionally induce expression of diphtheria toxin (DT) receptor (DTR) in GFAPcre+ cells. (b) Experimental timeline illustrating the sequence of behavioral testing on GFAPcre+ mice infused in the PFC with the AAV5-GFP-DIOCMV-DTRflag: sucrose test (ST), elevated plus maze (EPM), open field (OF), novelty suppressed feeding (NSF), locomotor activity (LM), novelty induced hypophagia (NIH), and forced-swim test (FST). Following DT administration (circle -0, square -0.1, triangle -5, and inverted triangle -20μg/kg), percent sucrose consumption in the ST was measured on day 1 (c), day 2 (d), day 3 (e) for 24 hours, and on day 4 (f) for 1 hour after 16hr fluid deprivation. Mice were also tested in the EPM (g), OF (h), NSF (i), and NIH (j). On day 8, mice percent sucrose consumption in the ST was measure (k) and were tested in the FST (l). Additional behavioral assessments (LM, water intake, and home-cage latency to feed and drink) can be found in Supplementary Table 2. Overall z-score of anxiety-like z-score (m) and anhedonia-like z-score (n) were calculated for each animal and group. Data are presented as individual animals and mean ± SEM. *\*p*< .05, **p< .01, and ***p<.001 as compared to DT0 group.

**Fig. 2:**
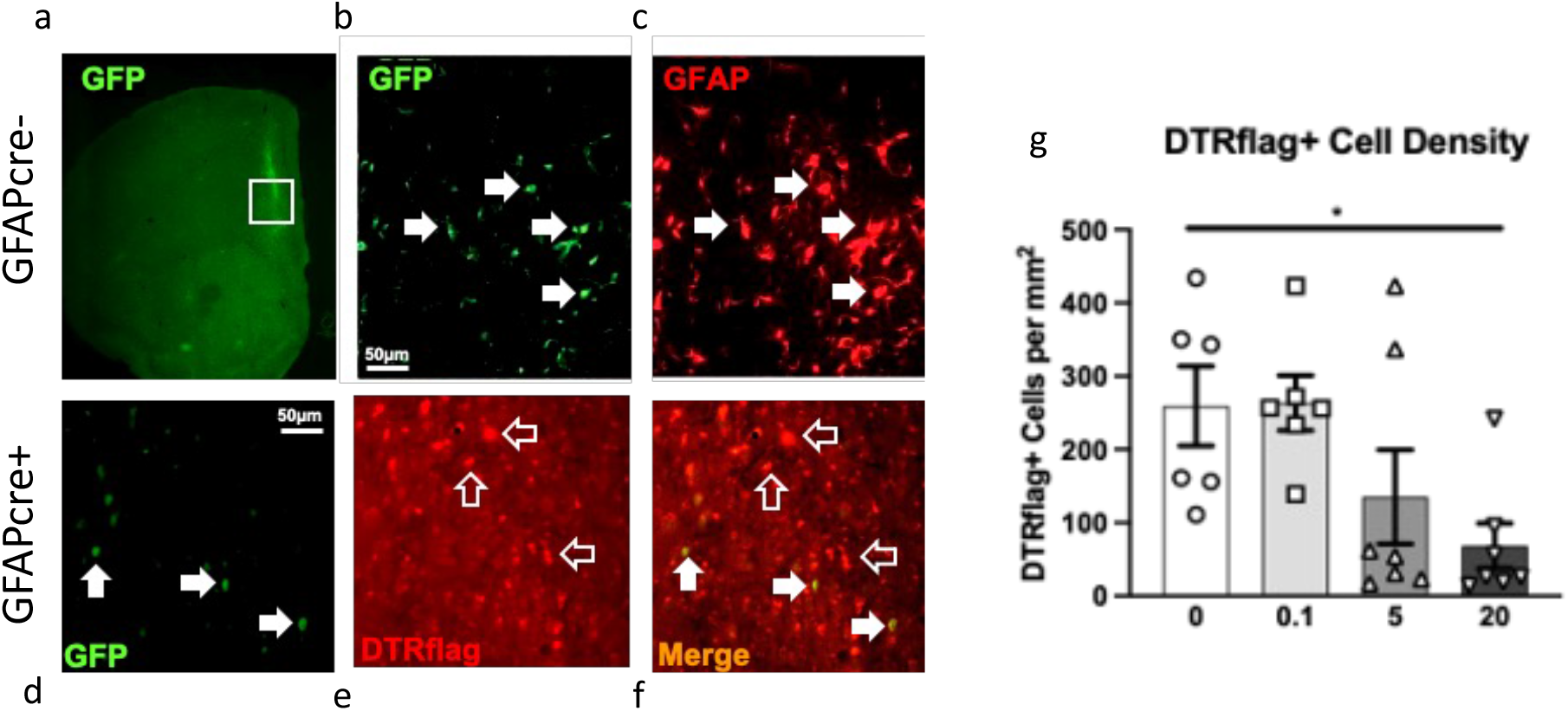
– Cell specificity of diphtheria toxin receptor (DTR) expression and cortical glial fibrillary acidic protein (GFAP)+ cell ablation. In GFAPcre-mice infused with AAV5-GFP-DIOCMV-DTRflag, the CMV promoter drives the expression of GFP as shown in a representative 2x image of GFP staining at the infection site (a). (b) High magnification of (a)-insert, with white arrows indicating GFP expressing cells which also express GFAP (c). In GFAPcre+ mice infused with AAV5-GFP-DIOCMV-DTRflag, the CMV promoter drives the expression of DTRflag. GFP and Flag immunostaining showed that some cells express very low levels of GFP (white arrow)(d), most infected cells express only DTRflag (empty arrow)(e), and cells expressing GFP also express flag (white arrow) (f). (b-f) Scale = 50um. (g) DTRflag cell density per mm2 was quantified in GFAPcre+ mice infused with AAV5-GFP-DIOCMV-DTRflag and treated or not with DT (circle -0, square -0.1, triangle -5, and inverted triangle -20μg/kg). Data are presented as individual animals and mean ± SEM. *\*p*< .05 as compared to DT0 group.

GFAPcre+ animals infused with AAV5-GFP-DIOCMV-DTRflag were tested in series of behavioral assays following DT or vehicle injections (Fig.1b). On day 1 of sucrose consumption testing, there was no significant effect of drug (F_(3,23)_ = 0.5, p>0.05; Fig.1c). On day 2, we found a significant effect of drug (F_(3,23)_=3.5, p<0.05), explained by a reduction in sucrose intake in the DT 20ug/kg group compared to vehicle (p<0.05, Fig.1d). On day 3, a similar significant effect of drug was found (F_(3,23)_=3.2, p<0.05) with both DT 5 and 20ug/kg groups, showing significantly reduced sucrose intake as compared to vehicle (p<0.05, Fig.1e). After 16hr fluid deprivation, sucrose intake for 1hr was measured on day 4. Results show a significant effect of drug (F_(3,22)_=6.9, p<0.01); post-hoc analysis revealed that sucrose intake in DT 5 and 20 ug/kg groups were significantly reduced as compared to vehicle (p<0.05; p<0.001; Fig.1f). Water consumption remained constant between groups (Table.S2). In the elevated plus maze (EPM), there was an effect of DT treatment on behavior (F_(3,22)_=3.3, p<0.05 Fig.1g), explained by a significant increase in time spent in the open arm in DT 0.1ug/kg as compared to vehicle; DT 5 ug/kg and 20 ug/kg did not differ from controls. In the open field (OF) and novelty-suppressed feeding test (NSF), there was no effect of DT treatment (OF: F_(3,22)_=2.1, Fig.1h; NSF: F_(3,22)_=1.9, Fig.1i). No change in home cage latency to eat was found between groups (Table S2). In the novelty induced hypophagia (NIH), there was a significant effect of drug (F_(3,22)_ = 3.2, p<0.05) explained by increased in the latency to drink in all 3 DT groups compared to the vehicle group (0.1ug/kg: p<0.01, 5ug/kg: p<0.05, 20ug/kg: p<0.05; Fig.1j). No change in home cage latency to drink was found between groups (Table S2). The effect of drug on percent sucrose consumption was confirmed on day 8 (F_(3,22)_=4.0, p<0.05), where 5 and 20ug/kg groups showed reductions in sucrose intake as compared to the vehicle group (p<0.05; Fig.1k). On day 8, animals were also tested in the forced swim test (FST), where we found no effect of drug on time spent immobile (2-6min: F_(3,22)_=1.9; 6-10min: F_(3,22)_=2.4; Fig.1l).

Analysis of anxiety z-score, calculated from average z-score of EPM, OF, and NSF performances, showed no effect of drug (F_(3,22)_= 1.6; Fig.1m). Analysis of the effect of drug on anhedonia z-score calculated by averaging z-scores in NIH on day 7 and sucrose consumption test on day 8, showed an effect of treatment (F_(3,22)_=5.8, p<0.01), explained by significant increase in anhedonia z-score in all DT groups as compared to the vehicle group (p<0.01; Fig.1n). There is no difference on locomotor (LM) activity between groups (Table S2).

In a control experiment following a similar design, the same AAV5-GFP-DIOCMV-DTRflag virus was infused into WT littermates. No significant effect of drug was found on sucrose intake (day 1:F_(3,24)_=1.0; day 2:F_(3,24)_ =0.2; day 3:F_(3,24)_=0.4), NSF (F_(3,24)_=0.3), NIH (F_(3,24)_=0.2), or in FST (2-6min:F_(3,24)_ = 1.4, p>0.05; 6-10min:F_(3,24)_=0.2, p>0.05) (Fig. S2). No difference in water consumption, latency to drink, or eat and LM activity were found (Table S2).

At the end of these behavioural assessments, animals were perfused, and we confirmed using immunohistochemistry the specificity of the virus employed. 70-80% of PFC GFAP+ astrocytes at the site of infection were colabelled with GFP in GFAPcre-mice were infused with AAV5-GFP-DIOCMV-DTRflag (Fig.2a-c). DTRflag+ cells were only detected in GFAPcre+ mice infused with AAV5-GFP-DIOCMV-DTRflag (Fig.2d-e). Quantification of DTRflag+ cell density revealed a significant effect of DT (F_(3,22)_=3.9, p<0.05) explained by a 73% reduction in DTRflag+ cells in the DT 20ug/kg group compared to vehicle treated animals (p<0.05; Fig.2g).

In two further cohorts we verified that the observed behavioral effects were specific to PFC GFAP+ cell depletion. First, we repeated the experiment described in Fig.1 using AAV5-GFP-mDIOCMV-DTRflag, which contains mutated lox-sites and thus leads to no DTR expression or cell depletion. There were no significant effects of DT on sucrose consumption (day 1:F_(1,10)_=0.2; day 2:F_(1,10)_=0.3; day 3:F_(1,10)_=1.9), NSF (F_(1,10)_=0.1), NIH (F_(1,10)_=0.2), or FST (2-6min:F_(1,10)_=1.0; 6-10min:F_(1,10)_=0.1, p>0.05) (Fig. S3). Water consumption, latency to drink, or eat and LM activity remained constant between groups (Table S2). Next, we infused the active virus, AAV5-GFP-DIOCMV-DTRflag, into the striatum of GFAPcre+ and WT mice and treated them with DT. In this experiment there was no main effect of genotype, drug, or drug*genotype interaction on sucrose consumption (day 1:genotype-F_(1,30)_=0.8, drug-F_(1,30)_=0.8, drug*genotype-F_(1,30)_=0.1; day 2:genotype-F_(1,30)_= 0.1, drug-F_(1,30)_=0.1, drug*genotype-F_(1,30)_=0.1; day 3:genotype-F_(1,30)_ = 0.1, drug-F_(1,30)_=0.1, genotype*drug-F_(1,30)_=1.4), in NIH test (genotype-F_(1,30)_=1.4, drug-F_(1,30)_=0.2, genotype*drug-F_(1,30)_=0.2), or in the NFS test (genotype-F_(1,30)_ = 0.1, drug-F_(1,30)_=0.6, genotype*drug-F_(1,30)_=1.4)(Fig. S4). Water consumption, latency to drink, or eat and LM activity remained constant between groups (Table S2).

### Enhancement of the activity of GFAP+ cells reverses the behavioral effects of chronic restraint stress(CRS)

We next performed a converse manipulation i.e. enhancing GFAP+ cell activity using a chemogenetic approach. To validate GFAP+ cell activation, we first conducted fiber photometry calcium imaging in animals infused in the PFC with AAV5-Zac2.1gfaABC1D-lck-GCaMP6f and AAV5-GFAP-hM3D(Gq)-mCherry to identify changes in GFAP+ calcium transients before and after administration of saline or clozapine-N-oxide (CNO). Repeated measures ANOVA of calcium transient peak frequency shows a main effect of drug (F_(1,9)_=8.3, p<0.05), no effect of time (F_(1,9)_ = 1.8), and a trend toward a drug*time interaction (F_(1,9)_=3.7, p=0.09). Further analysis shows a significant increase in peak frequency when mice were injected with CNO compared to baseline (F_(1,12)_=8.2, p<0.05), but not when they received saline (F_(1,6)_=0.2, p>0.05) (Fig.3c-d). At the end of the experiment, mice were sacrificed, and the expression and cell specificity of AAV5-Zac2.1 gfaABC1D-lck-GCaMP6f and AAV5-GFAP-hM3D(Gq)-mCherry were confirmed (Fig.3a-b).

**Fig. 3:**
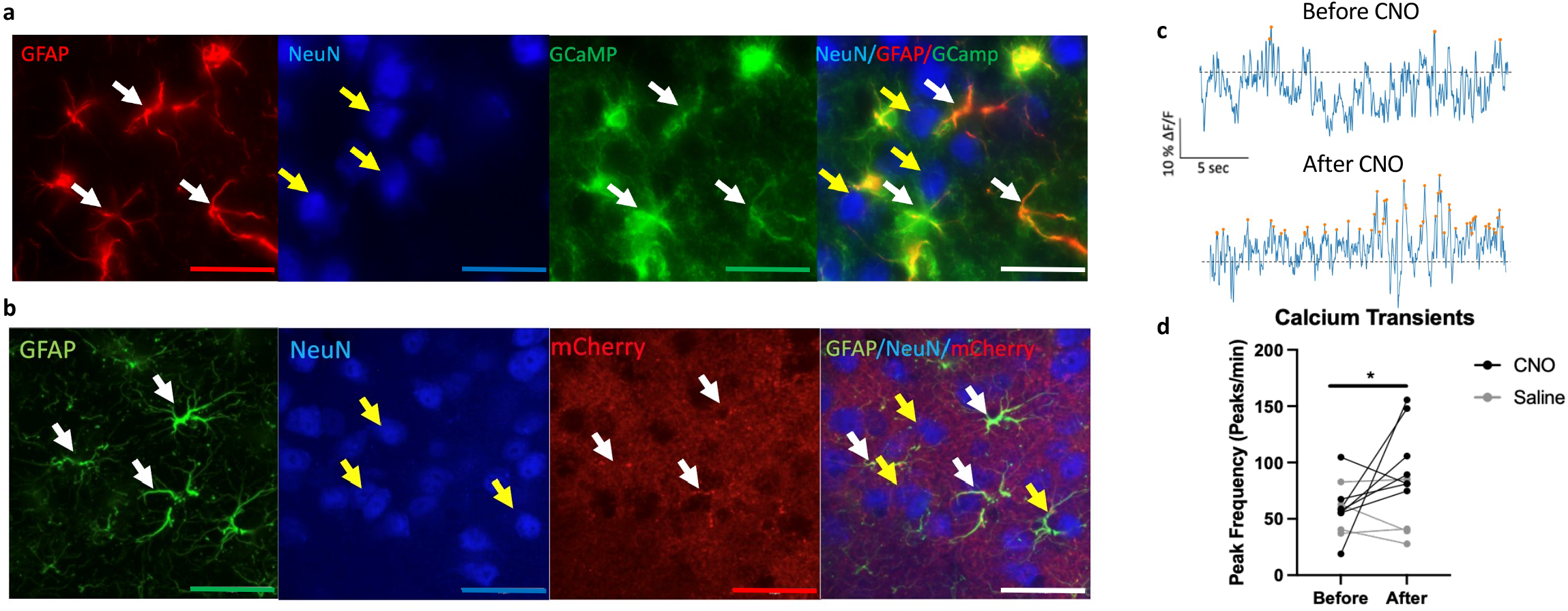
Virus cell specificity and enhancement of activity in PFC GFAP+ cells infused with AAV5-GFAP-hM3D(Gq)-mCherry virus. (a) Immunohistochemistry confirming AAV5-Zac2.1 gfaABC1D-lck-GCaMP6f virus specificity in PFC GFAP+ cells. White arrows indicate infected cells express GCaMP (green) and are co-labeled with GFAP+ cells (red). Yellow arrows indicate that NeuN+ cells (blue) are not co-labeled with either GFAP or GCaMP. Scale = 20um (b) Immunohistochemistry confirming AAV5-GFAP-hM3D(Gq)-mCherry virus specificity in PFC GFAP+ cells. White arrows indicate infected cells express mCherry (red) and are co-labeled with GFAP+ cells (green). Yellow arrows indicate that NeuN+ cells (blue) are not co-labeled with either GFAP or mCherry. Scale = 20um (c) Representative 30 second trace recording of calcium transients before and after acute CNO administration. Dotted black line: Z-score = 0. Orange dots indicates local maxima peaks that are 3 median absolute deviations (MAD) above the median. Y axis scale represents 10% z-score deltaF/F and X axis scale represents 5 seconds. (d) Peak frequency of calcium transients recorded for 30mins before and after acute administration of saline or clozapine-n-oxide (CNO) in animal infused with both viruses. Data represents peak frequency for individual recording for each animal before and after injection. *\*p*< .05 compared to before injection conditions.

We used this approach to test whether enhancing GFAP+ cell activity could reverse the behavioral effects of chronic restraint stress (CRS). We assessed sucrose consumption, coat state (CS), residual avoidance (RA) after the light challenge, and hourly time spent in the shelter in the PhenoTyper test in animals infused in the PFC with AAV5-GFAP-hM3D(Gq)-mCherry, with or without CRS for 6 weeks and with or without daily CNO treatment for the final four weeks (Fig.4a). Testing was performed weekly; here we describe the data obtained on week 2, before the start of CNO treatment, and in the final week of CNO treatment. Data from the other weeks can be found in the *Supplementary Results* and *Fig*.*S5*.

**Fig. 4:**
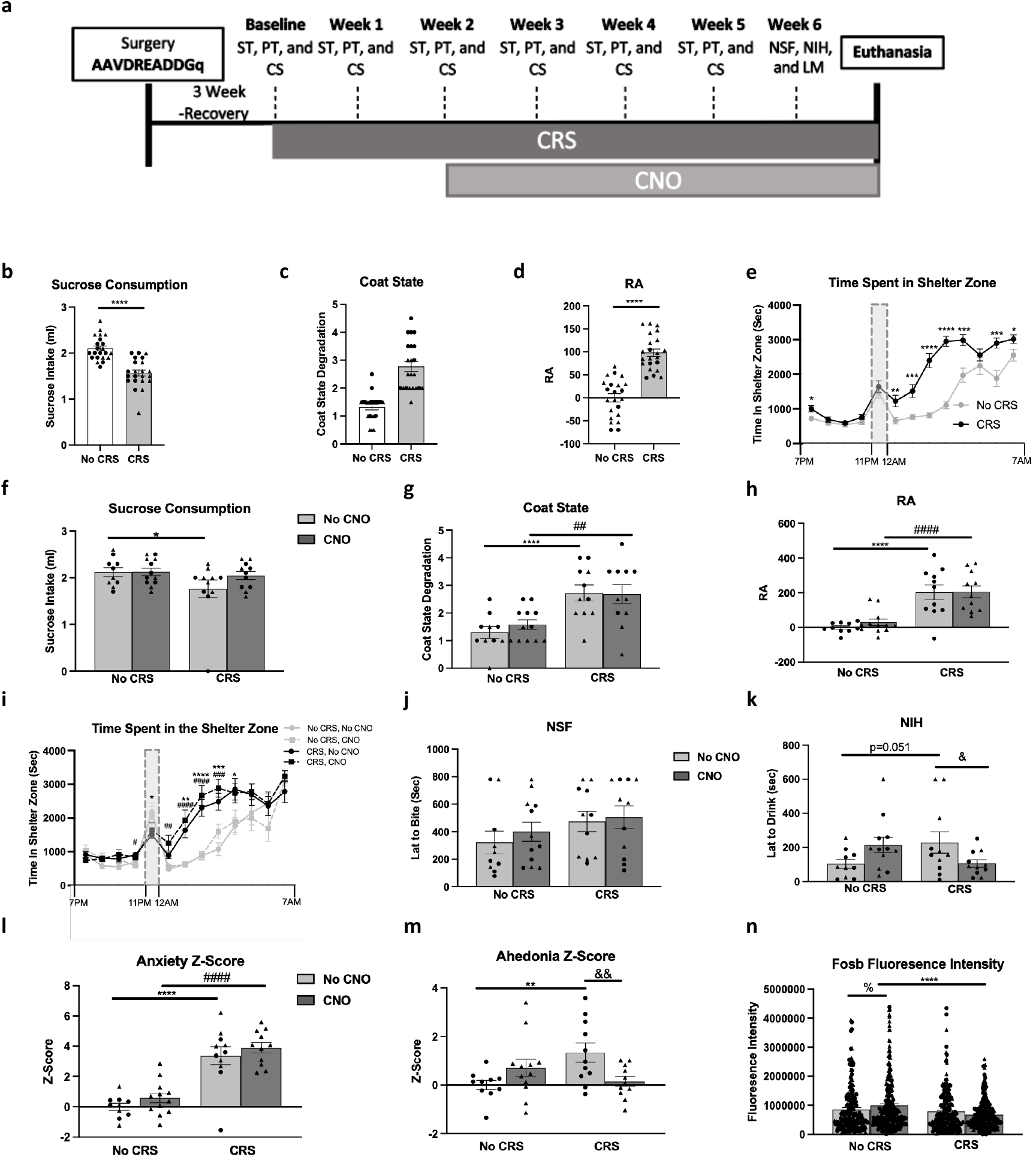
Cortical Glial Fibrillary Acidic Protein (GFAP)+ cell activity enhancement reverses chronic restraint stress (CRS)-induced anhedonia-but not anxiety-like behavior. (a) Experimental timeline illustrating the sequence of behavioral testing on mice infused in the PFC with the AAV5-GFAP-hM3D(Gq)-mCherry virus and subjected to CRS or not and treated with clozapine-n-oxide (CNO) or not: sucrose test (ST), PhenoTyper test (PT), coat state (CS), novelty suppressed feeding (NSF), novelty induced hypophagia (NIH), and locomotor activity (LM). (b-e) Behavioral assessment of mice subjected to 2 weeks of CRS or no CRS on sucrose consumption (b), coat state (c), residual avoidance (RA) (d), and hourly time spent in the shelter zone recorded in the PT (e). (f-k) Behavioral assessment of mice subjected to 5 weeks of CRS or no CRS and treated or not with CNO for the last 3 weeks of CRS on sucrose consumption (f), coat state (g), residual avoidance (RA) (h), and hourly time spent in the shelter zone recorded in the PT (i), NSF (j), and NIH (k). Overall z-score of anxiety-like z-score (l) and anhedonia-like z-score (m) were calculated for each animal and group. (n) Fluorescence intensity of Fosb per cell was quantified in 10 GFAP+/mCherry+ cells per hemisphere per animal. Females are shown as triangles and males as circles. Data are presented as mean ± SEM. Between group significant differences are marked as *\*p*< .05, **p< .01, ***p<.001, and ****p<.0001: No CRS+No CNO vs. CRS+No CNO; #*p*< .05, ##p< .01, ###p<.001,and ####p<.0001: No CRS+CNO vs. CRS+CNO; &*p*< .05, &p< .01, &p<.001, and &p<.0001: CRS+No CNO vs. CRS+CNO; %*p*< .05, %p< .01, %p<.001, and %p<.0001: No CRS+No CNO vs. No CRS+CNO.

As expected, 2 weeks of CRS induced anxiety and anhedonia (*20, 33*): mice exposed to CRS show a significant reduction in sucrose consumption (t_(1,42)_=37.9, p<0.0001; Fig.4b), no change in water consumption (Table S2), increase in CS degradation (t_(1,42)_=47.4, p<0.0001; Fig.4c), and increased RA (t_(1,42)_=66.7, p<0.0001; Fig.4d) compared to the no-CRS group. The RA was calculated from hourly recordings of time in the shelter zone following the light challenge, where we found a significant main effect of time (F_(1,42)_=90.8, p<0.0001), stress (F_(1,42)_=65.5, p <0.0001), and time* stress interaction (F_(1,12)_=8.7, p< 0.0001), explained by CRS mice spending more time in the shelter zone after the light challenge (p<0.0001;Fig.4e).

Analysis of the same behavioral tests following 3 weeks of CNO show a trend towards a main effect of stress in sucrose consumption (F_(1,40)_=3.3, p=0.07), but no main effect of drug (F_(1,40)_=1.4), or stress*drug interaction (F_(1,40)_=1.3). Post-hoc analysis revealed only a significant reduction in sucrose intake in the CRS+No CNO group compared to the No CRS+No CNO group (p<0.05; Fig.4f). No difference between groups was found on water consumption (Table S2). CS analysis revealed a main effect of stress (F_(1,40)_=23.0, p <0.0001), but no effect of drug (F_(1,40)_=0.2), or stress*drug interaction (F_(1,40)_=0.3), explained by a significant increase in CS degradation when comparing the 2 home cage control groups to their respective CRS groups (p<0.001;Fig.4g). ANOVA of RA revealed a main effect of stress (F_(1,40)_=40.9, p<0.0001), but no effect of drug (F_(1,40)_=0.3), or drug*stress interaction (F_(1,40)_=0.2). These effects are explained by a significant increase in RA when comparing the two home cage control groups to their respective CRS groups (p<0.05;Fig.4h). Specifically, mice exposed to CRS spend more time in the shelter zone after the light challenge, since hourly recordings of time in the shelter zone showed a significant main effect of stress (F_(1,40)_=28.8, p <0.0001), time (F_(1,40)_=58.5, p<0.0001), and stress*time interaction (F_(1,12)_=8.7, p<0.0001). However, groups receiving CNO were not different from their respective control groups since no effect of drug (F_(1,40)_=0.7), stress*drug interaction (F_(1,12)_=0.6), or drug*time interaction (F_(1,12)_=0.6) was found (Fig.4i).

During the final week of behavioral testing, in the NSF, there was no main effect of stress (F_(1,40)_=2.8), drug (F_(1,40)_=0.5) or stress*drug interaction (F_(1,40)_=0.1;Fig.4j). In the NIH, there was no main effect of stress (F_(1,40)_=0.1), or drug (F_(1,40)_=0.1), but there was a drug*stress interaction (F_(1,40)_=7.4, p<0.01) reflecting a significant increase in latency to drink in the CRS+No CNO group compared to No CRS+No CNO (p<0.05) and reversal in CRS animals receiving CNO (p<0.05;Fig.4k). tested the potential contribution of sex as a factor in the aforementioned analyses Analysis of anxiety z-score calculated from average z-score of week 5 RA and NSF performances show a main effect of stress (F_(1,40)_=69.55, p<0.0001), but no effect of drug (F_(1,40)_=1.9), or stress*drug interaction (F_(1,40)_=0.1), explained by a significant increase in anxiety z-score when comparing the 2 home cage control groups to their respective CRS groups (p<0.0001;Fig.4l). Analysis of the effect of drug on anhedonia z-score calculated by averaging z-scores in week 5 sucrose consumption and NIH showed no main effect of stress (F_(1,40)_=1.5), or drug (F_(1,40)_=0.6), but a stress*drug interaction (F_(1,40)_=9.3, p<0.01), explained by a significant increase in anhedonia z-score in the CRS+No CNO group compared to the No CRS+No CNO group (p<0.01) and reversal of this effect in CRS+CNO group (p<0.01;Fig.4m). We also established that administration of CNO had no effects on the various behavioral readouts used in this study by treating CRS or home cage control mice with CNO for 3 weeks (Supplementary Results and Fig. S6).

To further confirm that GFAP+ cell activity is enhanced following administration of CNO, we analyzed Fosb fluorescence intensity in mCherry+/GFAP+ cells. Analysis of Fosb fluorescence intensity revealed a main effect of stress (F_(1,860)_=14.9, p<0.0001), no effect of drug (F_(1,860)_=0.1, p>0.05), but a stress*drug interaction (F_(1,860)_=6.1, p<0.05). Post-hoc analysis showed a significant increase in Fosb fluorescence intensity in No CRS+CNO group as compared to No CRS+No CNO (p<0.05), as well as decrease in CRS+CNO group compared to No CRS+CNO (p<0.05;Fig.4n).

## DISCUSSION

In this study, we investigated the relationship of PFC GFAP+ cells to depressive-like behaviors in mice. We found that mice with PFC GFAP+ cell depletion show reductions in sucrose intake, within 2 days after DT administration and lasting up to 8 days, as well as an increase latency to drink in NIH. This suggests that PFC GFAP+ astroglia depletion induced anhedonia-like behavioral deficits, which was confirmed in the analysis of overall anhedonia z-score. However, mice with GFAP+ cell depletion showed no behavioral deficits in the FST or in any of the independent tests for anxiety-like behaviors (EPM, OF, NSF) or overall anxiety z-score. These effects were specific to the GFAP+ cell depletion and not due to side effects of DT, since no difference between groups were found in WT littermates or when using a mutated virus. In addition, the behavioral effects of PFC GFAP+ cell depletion were region specific since no anhedonia-like behavioral deficits were observed following striatal GFAP+ cell depletion. Overall, our results suggest that cortical GFAP+ cell depletion induce anhedonia-like but not anxiety- or helplessness-like deficits.

In a converse experiment, we tested the effects of chemogenetic activation of GFAP+ cells in the mPFC. After confirming that 2 weeks of CRS reduced sucrose intake and increased coat state degradation and RA, we used an hM3D(Gq) receptor to enhance PFC GFAP+ cell activity. We showed that enhancing PFC GFAP+ cell activity for 3-4 weeks reversed CRS-induced deficits in sucrose intake and latency to drink in the NIH. Overall anhedonia z-score was increased by CRS and was reversed by enhancing PFC GFAP+ cell activity. However, enhancing PFC GFAP+ cell activity was unable to reverse CRS-induced anxiety-like effects measured independently in the PhenoTyper and the NSF test or overall, on anxiety z-score. The increase in PFC GFAP+ cell activity was confirmed following CNO administration using calcium imaging (acutely) and Fosb immunohistochemistry (chronically). Altogether, we demonstrated that enhancing PFC GFAP+ cell activity reversed CRS-induced anhedonia-but not anxiety-like behavior.

Viruses expressing DTR have been previously employed to induce cell and/or region-specific cell apoptosis upon injection of DT (*34-37*). In this study, we observed reductions of GFAP+ cells, *in vitro* using astroglial culture and a cell survival assay, and *in vivo* by quantifying the number of DTRflag positive cells following DT administration. This is consistent with previous studies using this approach to target and ablate specific neuronal populations within select brain regions (*34, 36*). The cytotoxic properties of DT are contingent upon DT binding to simian DTR, which results in termination of protein synthesis and eventually apoptosis (*35*). Some studies have suggested that DT may cause locomotor side effects in WT mice (*38*), however, we found no change in locomotion following DT treatment in WT or GFAPcre+ mice. Behavioral effects of PFC glial depletion have been previously investigated using non-specific gliotoxins, which induce depressive-like deficits (*21*), increase alcohol preference (*22*), and impair cognitive flexibility (*23*).

However, given the plethora of evidence reporting that astroglia reductions in the PFC of MDD post-mortem and rodent chronic stress models seem to affect preferentially the GFAP+ cell population (*8,28, 29, 39, 40*), we felt necessary to explore the specific contribution of GFAP+ cell pathology in the development of depressive-like behavior. Interestingly, the behavioural effects of PFC GFAP+ cell depletion on anhedonia-like behavior appeared as early as two days following DT; such effects are typically observed only after several weeks of chronic stress (*20, 41*). This suggests that in animals exposed to chronic stress, the observed cortical astroglia loss may be a turning point for the apparition of the anhedonia-like deficits. This study emphasizes a role of PFC GFAP+ cell loss or dysfunction in the development or onset of anhedonia-like behavior.

CRS is widely used to model chronic stress and is known to induce anhedonia- and anxiety-like behavior, physical and cellular deficits in rodents (*20, 33, 42-44*). We previously demonstrated that 2 weeks of CRS increased coat state deterioration (*20, 44, 45*), decreased sucrose consumption (*20, 33*), and increased RA (*33, 45*). These effects last 5-6 weeks (*20, 33, 44, 45*). CRS behavioral deficits are similar to those observed with chronic unpredictable (mild) stress or social defeat models (*44-47*). Here, we opted to use the CRS model as we previously demonstrated that CRS induced GFAP+ cell atrophy (*20*) but knowing that CRS animals display temporary locomotor hyperactivity upon handling, rendering short tests such as EPM, OF, and FST unusable in this model (*45, 48*).

To enhance astroglial function, cells were infected with a virus expressing hM3D(Gq) receptor which is activated exclusively via the designer ligand CNO (*49, 50*). It was previously shown the activation of this DREADD receptor in astrocytes and the resulting calcium mobilization regulate synaptic processing and plasticity (*24, 51*). Gi-coupled DREADD receptors have been used to decrease activity in neurons but were shown to have the opposite effect in astrocytes (*24*). For this reason, we could not use the Gi method for reducing GFAP+ cell function and instead opted for the astroglial depletion approach, which more closely models the GFAP+ astroglial density reductions associated with MDD pathology (*7, 8, 17, 18, 28*). Previous studies have successfully demonstrated in several brain regions that activation of the Gq-DREADD receptor in GFAP+ astroglia enhances cell activity (*52-55*). Here we used fiberphotometry to confirm increased cortical GFAP+ calcium flux following stimulation of the Gq-DREADD receptor in infected cells upon application of CNO. Since CNO can be metabolized into a clozapine-like substances (*56, 57*) and clozapine can have potential anti-stress properties, concerns were raised regarding the use of chronic CNO in studies investigating emotion related-behavior (*58*). Therefore, we verified using the conventional treatment of chronic CNO in drinking water (*59*) that CNO treatment could not reverse or prevent CRS effects in the relevant behaviors and time periods of this study.

We also confirmed PFC GFAP+ cell activation chronically using Fosb fluorescence intensity. The Fosb family includes Fosb and ΔFosb proteins, both recognized by the antibody employed in this study. ΔFosb increases in neurons following chronic stress or cell activation (*60, 61*). The use of Fosb labelling for detecting astrocyte activation is limited and was shown to remain unchanged after chronic stress, which we confirmed here (*60, 62*). We corroborated PFC GFAP+ enhanced activity using the DREADD approach (increases GFAP+/Fosb+ cells in no CRS animals), which rules out hM3D(Gq) receptor desensitization following chronic CNO treatment in our experiment (*50*). However, it remains unclear why this increase in Fosb+ cells is blunted in CRS animals despite GFAP+ cell activity enhancement reversing the CRS-induced behavioral effects. The mechanisms involved in this divergence may need to be investigated in future studies.

In this study, we demonstrated that PFC GFAP+ cell depletion induces anhedonia-like behavior and that enhancing PFC GFAP+ cell activity reversed CRS-induced anhedonia-like deficits. Both manipulations had little or no effect on anxiety-like behavior. This could be due to the PFC regional specificity of GFAP+ cell targeting since chemogenetic activation of GFAP+ cell activity in the amygdala reduced fear expression, altered fear memory and modulated PFC-amygdala communication (*52, 63*). Another study has also pointed out a role of hippocampal astrocytes in anxiety-like behaviors, specifically demonstrating that activated hippocampal astrocytes induced anxiolytic-like effects (*64*). Here, we establish a specific connection between PFC GFAP+ astroglia number or activity and the modulation of anhedonia-like behavior. We can therefore speculate as to the mechanisms through which cortical astrocytes regulate behavior. Since GFAP+ cells main functions are to modulate neuronal and synaptic function, it is possible that the regulation of behavior occurs through these processes. Astrocytes regulate neuronal activity through the uptake of neurotransmitters released from presynaptic terminals (*24, 51*), release of gliotransmitters and neurotrophic factors (*16, 65*), in addition to the regulation of synapse formation, maintenance and plasticity (*12, 16, 66*). These mechanisms and functions are affected by stress and are thought to be involved in MDD pathology (*7, 67*). Therefore, PFC GFAP+ cell depletion would likely result in alterations of the function of the pyramidal neurons and interneurons within the cortical circuit and impairment of synaptic transmission or integrity. Such changes and PFC GFAP+ cell reductions have been observed following chronic stress in rodents (*8, 68, 69*). Conversely, we speculate that enhancing PFC GFAP+ cell activity in stressed animals partially prevents of these neuronal alterations and reverse the behavioral effects of chronic stress. However, to test these hypotheses, future studies would need to focus on the consequences of manipulation of GFAP+ cell number or activity on neuronal and synaptic function.

This study is not without its limitations. First, these experiments were performed with a relatively small number of animals. However, even with a limited number, we were able to detect significant effects both cellularly and behaviorally following GFAP+ cell depletion and activity enhancement. Another limitation is that the depletion studies were conducted only in males; this was necessary as we used the GFAPcre+ females for breeding to avoid germline recombination and for generating pups for the *in vitro* work. In the calcium imaging study we chose to characterize the effects of GFAP+ cell activity enhancement acutely. In a pilot experiment, we attempted repeated calcium recordings following chronic CNO administration. However, within animal baseline variability in calcium transients and weekly fluctuations in the vehicle group hindered our ability to confidently assess chronic CNO treatment effects using fiber photometry. We instead opted to use Fosb fluorescence intensity to confirm chronic enhancement of GFAP+ cell activity. Although we did successfully validate GFAP+ cell activity enhancement using this approach, it is important to mention that we encountered difficulties when quantifying Fosb fluorescence intensity as its cellular localization differs from that of the GFAP and mCherry staining which were used for tracing astrocyte processes. This may have caused smaller effect sizes and greater group variability potentially impeding the detection of increases in GFAP+ activity in animals subjected to stress who display GFAP+ cell atrophy (*20, 30*). Finally, in this study we chose to focus on the behavioral effects associated with astroglial depletion and activity enhancement; the question remains as to the neuronal consequences of such manipulations.

In summary, our study demonstrates that PFC GFAP+ astrocytes play a causal role in modulating anhedonia-like behavior in mice. We found anhedonia-like effects following PFC GFAP+ cell depletion and oppositely reversal of chronic stress-induced anhedonia-like deficits after GFAP+ cell activity enhancement in the PFC. This work further highlights the importance of astrocyte dysfunction in the development of the behavioral deficits associated with stress-related illnesses and highlights its potential as a target for antidepressant treatment.

## METHODS

### Animals

C57BL/6.Cg-Tg(GFAP-cre)77.6Mvs/2J transgenic mouse line (Jackson Laboratories, stock #024098) were bred in-house. GFAP-cre females were used for breeding and for generating pups for *in vitro* development of the GFP-DIOCMV-DTRflag plasmid. Male 8-12-week-old GFAP-cre and Wild-type (WT) mice littermates were used for the depletion experiments. WT mice (C57/Bl-6, Jackson Laboratories, lot # 000664, 50% female), approximately 8-9 weeks old, were used for the astrocyte activity enhancement studies.

Mice were single housed under normal condition at a consistent temperature (∼25°C) with 12-hour light/ dark cycle and ad libitum access to food and water (except for deprivation of food or water for behavioral testing). All procedures were in accordance with the Yale University Care and Use of laboratory animals (YACUC) guidelines and the Canadian Council on animal care further approved by Centre for Addiction and Mental Health (CAMH) animal care committee (CAMH-ACC).

### Virus Information

We adapted the construct from (*36*) by strategically placing the loxP and lox2722 sites to flank the CMV promoter (Fig.1a). The construct is designed so that in non-cre cells CMV drives the expression of GFP and in cre cells CMV drives the expression of the DTR fused to a FLAG epitope for ready immunohistochemical identification. This was confirmed by transient transfection in primary astrocyte cultures generated from GFAPcre+ pups (*supplementary methods*), as well cell depletion upon application of DT (Fig. S1). The control virus is identical except that the lox sites are mutated to prevent recombination. The constructs were packaged into AAV5 in HEK293 cells and infused into the PFC. For the fiberphotometry and GFAP+ cell activity enhancement experiments, commercially available viruses were used: pAAV-GFAP-hM3D(Gq)-mCherry (Addgene virus #50478) and pZac2.1 gfaABC1D-lck-GCaMP6f virus (Addgene virus #52924).

### Chronic Restraint Stress (CRS) Procedure

The CRS procedure was performed as in (*20, 33, 44, 45*) *(supplementary methods)*.

### Surgery and Drug Protocol

GFAP+ cell depletion: GFAP-cre mice (n=6-7/group) and WT littermates (n=8/group) were infused in the PFC with the AAV5-GFP-DIOCMV-DTRflag (coordinates: AP+ 2, DL −/+0.5, Depth −3 from Bregma). In a separate cohort, GFAP-cre mice (n=7/group) were infused in the PFC with a AAV5-GFP-mDIOCMV-DTRflag. Additionally, to evaluate region specificity of the behavioral effects, GFAP-cre (n=7/group) mice were injected in the striatum with 0.5ul of AAV5-GFP-DIOCMV-DTRflag (coordinates:AP+1.5, DL−/+1.5, Depth−3; Titer≥7×10^12^vg/mL). After a 3-week recovery period, mice were injected with DT (0.1, 5, or 20ug/kg, i.p) or saline every evening before the sucrose test for the first 3 days at 6pm. The animals were then tested every following day in behavioral assays (Fig.1b).

PFC GFAP+ cell activity enhancement fiberphotometry: C57Bl/6 mice (n=8, 50% females) were infused in the PFC (50% right hemisphere) with 0.5ul of AAV5-GFAP-hM3D(Gq)-mCherry (Titer≥7×10^12^vg/mL) and 0.5ul of a AAV5-Zac2.1 gfaABC1D-lck-GCaMP6f (Titer≥7×10^12^vg/mL). Fiberoptic cannula (Doric Lenses Inc., Code: MFC_400/430-0.66_5mm _MF2.5_FLT) was implanted in the same hemisphere. After 3-week recovery, mice were given injections of clozapine-N-oxide (CNO) dissolved in saline (5mg/kg, i.p) or saline.

PFC GFAP+ cell activity enhancement: C57Bl/6 mice (n=12/group, 4 groups, 50% female) were bilaterally infused in the PFC with 0.5ul of the AAV5-GFAP-hM3D(Gq)-mCherry. Upon 3-week recovery and baseline behavioral assessment, half of the animals were subjected to CRS for 2 weeks for induction of depressive-like deficits. Groups were then split and were given a 5mg/kg daily dose of CNO dissolved in their drinking water or normal water. Animals were behaviorally assessed weekly, with additional behavioral assays performed in the last week and were euthanized (Fig.3a). We also conducted a control study assessing the behavioral effects of CNO in control, non-surgerized mice subjected to or not to CRS (C57Bl/6 mice, n=10/group, 4 groups, 50% female) (*supplementary method*).

### Behavioral Assessments

For the PFC GFAP+ cell abalation experiment daily sucrose consumption was measured for 3 days and percent sucrose consumption was calculated from the saline controls. Animals were then tested everyday in the 1-hour sucrose consumption test, EPM, OF, NSF, NIH, LM activity, retested in daily percent sucrose consumption on day 8 and then assessed in the FST (Fig.1b). For the PFC GFAP+ cell activity enhancement study, mice were tested weekly for 5 weeks in the 1-hour sucrose consumption test, time spent in the shelter in the PhenoTyper test, and CS. On week 6, performances in the NIH, NSF, and LM activity were also measured (Fig.3a).

The NSF, NIH, EPM, OF, FST and LM activity tests were performed as in (*44, 70-72*). For the stress study, the 1-hour sucrose consumption test, PhenoTyper Test, LM activity, and CS were performed as in (*20, 33, 44, 45*). Anxiety and anhedonia z-scores were calculated as in (*44, 70*). *See supplementary methods section*.

### Fiberphotometry Recording and Analysis

Fiberphotometry was performed using the Doric Studios (Quebec, CA) fiber photometry console with the 406 isosbestic channel and the 465 green channel with a sampling rate of 12kS/sec. Prior to recording, animals were placed into the recording room for a 1hr habituation and then recorded for 30 mins for baseline assessment. Mice were then injected with saline or CNO and subsequently recorded for 1-hour. Given that the peak of CNO activation is expected within 30 mins post injection, we focused our analysis on that window (*24, 57*). After recording, the raw data was exported, down sampled to 1 kS/sec, detrended to correct for photobleaching using the airPLS method (*73*). In short, the analysis was conducted using the GuPPy program, where the excitation signal was regressed against the isosbestic signal and deltaF/F was calculated separately for the 30mins before and after the CNO or saline administration. These periods were then normalized by z-score to standardized signals. For each recording, high amplitude events (greater then 2 median absolute deviations (MADs) above the median) were filtered out. Peaks with local maxima greater than the threshold (set at 3 MADs above the median) were quantified, summated, and divided by the duration of each recording to calculate peak frequency (peaks/min) (*74*).

### Immunochemistry, imaging, and quantification

Animals were perfused under anesthesia as in (*20*). Brain and tissue preparation, and immunochemistry protocols with employed primary/secondary antibodies can be found in the *supplement material and Table S1*. Validation of the construct, viruses, and quantification of the number of Flag+ cells and Fosb intensity are described in the *supplementary materials*.

### Statistical Analysis

For the depletion, fiberphotometry, and enhancement experiments, 9 animals total were removed due to attrition from surgery/cannula loss, mislocalization of the virus/cannula, or adverse reactions to stress. We removed one outlier based on its behavioral performances in multiple tests (2 standard deviations above or below the mean). Only the remaining animals were used for the analysis and in the results section. For analysis of the Fosb intensity, 16 out of 880 cells were outliers as they displayed intensities of more than 3 standard deviations above or below the group mean. Statistical analysis was performed using StatView software 5.0 (Berkley, CA, USA) and graphs were generated using Prism 9 (San Diego, CA, USA). ANOVAs were used to determine the main effects of stress and/or drug in single read-out tests with 3 or more groups. If the number of groups was 2, a t-test was performed. We used repeated-measures ANOVAs for the longitudinal data. Fisher’s test was used for post-hoc analysis.

## Supporting information

Fig.S1, Fig.S2, Fig.S3, Fig.S4, Fig.S5, Fig.S6, Fig.S7, Table.S1, Table.S2, Table.S3 will link to SuppFigures.pdf

Fig.S1, Fig.S2, Fig.S3, Fig.S4, Fig.S5, Fig.S6, Fig.S7, Table.S1, Table.S2, Table.S3 will link to this file

## ACKNOWLEDGEMENTS

We would like to acknowledge the contributions of the Yale and CAMH animal facility personnel for animal care and genotyping services, to include Lori Dixon, Kristen Fournier, Katrina Deverell, Shelia Ambata and Leila Tick. We would also like to thank Dr. Engelhardt at the University of Iowa for providing AAV5 helper plasmids.

This work was supported by the Brain and Behavior research foundation (NARSAD, PI: MB), National Institute of Mental Health (NIH-NIMH R01 MH081211, PI: GS), the CAMH discovery fund (PI: M.B.), the Canadian Institutes of Health Research (PGT165852, PI: M.B.) and the Campbell Family Mental Health Research Institute. JM was supported by the CIHR Doctoral Fellowship (201810GSD-4221 05-DRA-CFAA-297096), YB by CIHR Post-doctoral Fellowship (202110MFE-472592-FPP-CEAH-93191), the CAMH Discovery Fund Fellowship (2021-0859) and the CAMH womenmind Fellowship (2021-0863).

## AUTHOR CONTRIBUTIONS

SAC, ES, RSD, CP, and MB designed the experiments and provided conceptual assistance regarding methods and techniques. MX, CP, SW, PL and MB performed DTR virus validation in culture. MB, AEL, and VD conducted the depletion behavioral experiments and SAC conducted their analysis. SAC and MC performed the Fosb imaging. SC and YB conducted the fiberphotometry experiments and analysis with guidance from JM and RCB. SAC conducted all the enhancing astroglial function behavioural experiment with their analyses. SAC and MB wrote the manuscript. ES, GS, CP and MB reviewed and edited the manuscript. Supervision and funding acquisition was provided by MB.

