## Supplementary material for "Prefrontal Cortex Astroglia Modulate Anhedonia-like Behavior": Fig.S1, Fig.S2, Fig.S3, Fig.S4, Fig.S5, Fig.S6, Fig.S7, Table.S1, Table.S2, Table.S3 will link to SuppFigures.pdf

**Word Count:** 4774
**Number of Supplementary Figures:** 7
**Number of Supplementary Tables:** 3
**Number of Supplementary References:** 15

### SUPPLEMENTARY METHODS

*Viral Construct:*

We adapted the viral construct used in (*1*). Specifically, we used a modified FLEX cassette (Fig. 1a) to invert cytomegalovirus (CMV) sequence in the presence of Cre-recombinase. The construct contained two inverted cre-recognition sites (loxP and lox2722) flanking CMV allowing for the CMV promoter, which drives the expression of enhanced green fluorescent protein (eGFP) in non-cre condition, to invert and induce expression of diphtheria toxin receptor (DTR) in the presence of cre-recombinase. Cre-expressing cells are thus made susceptible to depletion by systemic administration of DT. The simian DTR gene is fused with the flu antigen (FLAG) epitope tag at its C-terminal was placed in the antisense orientation before the FLEX cassette. The control vector we used contained a similar sequence but with multiple point mutations in the 3′ loxP and lox 2722 sites, preventing inversion of CMV and allowing for the expression of eGFP (never DTR) regardless of Cre-expression. The mutated sequence can be found in (*1*). The sequences were cloned into an adenoassociated virus (AAV)2-1 backbone (AAV cis) and recombinant AA5 viral stocks were generated as previously described in (*1, 2*). Briefly, HEK293 were transfected with the AAV cis, R2 and T5 helper plasmids (4ul/ml/plasmid; T5 was generously provided by Dr. Engelhardt at the University of Iowa) using transfection reagent of Lipofectamine (Life Technologies). 3 days after transfection, cells pellets were collected and then lysed in freeze/thaw cycles in dry ice and ethanol. Lysates were filtered using amicon® and concentrated using the ultra-15 centrifugal filter units (Millipore). Using real-time PCR, we found that the final titer was 5-7 $\times 10^{12}$ vg/mL.

*Primary Astrocyte Culture:*

P0 mice were generated from GFAPcre-/+ females and WT males. Each prefrontal cortex (PFC) were dissected manually and astrocyte cultures were prepared as in (*3*). Tails were collected for genotyping. Cells were mechanically dissociated (20 passages through a 2mm diameter needle) and cell suspensions were diluted in 12ml of DMEM + fetal calf serum, 4 mM glutamine, 50 U/mL penicillin and 50 lg/mL streptomycin for each pup. Cells were seeded on 6 well plates all without coating (2 mL per well), but half of the wells contained coverslips to allow for immunocytochemistry. They were maintained in incubators at 37°C in a humidified atmosphere of 5% $CO_2$/95% air. Medium was changed weekly until confluence is reached (12–14 days). Astrocyte were then differentiated in defined medium containing DMEM supplemented with 4 mM glutamine, 5 lg/ mL insulin, 0.5 mg/mL bovine serum albumin fatty acid free, 50 U/mL penicillin, 50 lg/mL streptomycin, and 0.25 mM dBcAMP. The dBcAMP-differentiated cultures comprised more than 95% of GFAP-positive cells. Following the genotyping protocol described for this mouse line (Jackson Laboratories, guidelines for stock #024098), each plate was marked according to the genotype of the pup used to generate the culture.

The constructs were then tested using transient lipofectamine transfection, following the manufacturer's instructions. 1μg DNA was added to 200μL Opti-MEM medium and 4μL of lipofectamine then applied to each well, mixed, and then incubated at room temperature for 6 hours. Media was then changed back to defined medium for 5 days. Cells from half of the plates were rinsed with PBS and fixed with 4% paraformaldehyde/PBS solution for 1 hour and kept in PBS until immunostaining using antibodies specified in (Table.S1). The other half of the cells were used for the cell survival assay. Diphtheria toxin (DT) (D0561, Sigma) was applied for one hour and MTT assay was performed as in (*3*).

*Surgeries and Drug Protocol:*

PFC GFAP+ cells depletion: GFAP-cre mice (n=6-7/group) and WT littermates (n=8/group) were infused in the PFC with the AAV5-GFP-DIOCMV-DTRflag (coordinates: AP+ 2, DL −/+0.5, Depth −3 from Bregma). In a separate cohort, GFAP-cre mice (n=7/group) were infused in the PFC with a AAV5-GFP-mDIOCMV-DTRflag. Additionally, to evaluate region specificity of the behavioral effects, GFAP-cre (n=7/group) mice were injected in the striatum with 0.5ul of AAV5-GFP-DIOCMV-DTRflag (coordinates:AP+1.5, DL−/+1.5, Depth−3; Titer≥$7\times10^{12}$vg/mL). After a 3-week recovery period, mice were injected with DT (0.1ug/kg, 5ug/kg, 20ug/kg, i.p) or saline every evening before the sucrose test for the first 3 days at 6pm. The animals were then tested every following day in behavioral assays (Fig.1b).

PFC GFAP+ cell activity enhancement fiberphotometry: C57Bl/6 mice (n=8) were infused in the PFC (50% right hemisphere) with 0.5ul of AAV5-GFAP-hM3D(Gq)-mCherry (Addgene virus # 50478; Titer≥$7\times10^{12}$vg/mL) Designer Receptor Exclusively Activated by Designer Drug (DREADD) virus and 0.5ul of a AAV5-Zac2.1 gfaABC1D-lck-GCaMP6f virus (Addgene virus #52924; Titer≥$7\times10^{12}$vg/mL). Fiberoptic cannula (Doric Lenses Inc., Code: MFC_400/430-0.66_5mm _MF2.5_FLT) was implanted in the same hemisphere. After recovery for 3 weeks, mice were given injections of clozapine-N-oxide (CNO) dissolved in saline (5mg/kg , i.p) or saline.

PFC GFAP+ cell activity enhancement: C57Bl/6 mice (n=12/group, 4 groups, 50% female) were bilaterally infused in the PFC with 0.5ul of the AAV5-GFAP-hM3D(Gq)-mCherry and left to recover for 3 weeks. Upon recovery and baseline behavioral assessment, half of the animals were subjected to CRS for 2 weeks for induction of depressive-like deficits. Groups were then split and were given a 5mg/kg daily dose of CNO dissolved in their drinking water or normal water. Animals were behaviorally assessed weekly, with additional behavioral assays performed in the last week and were euthanized (Fig.3a).

CNO effect in CRS animals: C57Bl/6 mice (n=10/group, 4 groups, 50% female) were assessed in baseline behavioral tests, half of the animals were then subjected to CRS for 5 weeks. These groups were also split in half and were given a 5mg/kg daily dose of CNO dissolved in their drinking water or normal water. Animals were behaviorally assessed weekly, with additional behavioral assays performed in the last week and were euthanized (Fig.S6).

*Chronic Restraint Stress (CRS) Procedure:*

The CRS procedure entails animals being placed in a falcon tube, with small hole on either end for sufficient air flow and a place for their tail, for 1 hr, twice a day approximately 1-2hrs apart and at least 1hr before behavioral testing. This CRS model has been consistent in providing both behavioral and cellular deficits associated with chronic stress in our labs previous work (*4-7*).

*Behavioral Assessments:*

For the GFAP+ cell depletion studies, the animals were tested in the sucrose consumption test, daily sucrose consumption was measured for 3 days and daily percent sucrose consumption was calculated from the saline controls. Animals were then tested everyday in the 1-hour sucrose consumption test, elevated plus maze (EPM), open field (OF), novelty supressed feeding (NSF), novelty induced hypophagia (NIH), locomotor (LM), retested in daily percent sucrose consumption on day 8 and then assessed in the forced swim test (FST) (Fig.1b).

For the PFC GFAP astroglial activity enhancement study, mice were tested weekly for 5 weeks in the one-hour sucrose consumption test, time spent in the shelter in the PhenoTyper test (PT), and coat state assessment (CS). On week 6, performances in the NIH, NSF, and LM were also measured (Fig.3a). EPM, OF, and FST were not performed in these studies since it was shown that CRS animals display temporary locomotor hyperactivity upon handling, rendering short tests such as EPM, OF, and FST unusable in this model (*5, 8*).

*Sucrose Consumption Testing (ST):* This test is used as a consistent measure of anhedonia-like behavior (*5-7, 9, 10*). All sucrose consumption tests for the GFAP+ cell depletion studies were done with 1% sucrose solution, while the GFAP+ cell activity enhancement studies were done with 2% sucrose solution. We opted for these conditions because mice subjected to CRS and surgerized do not drink sufficient sucrose at 1% concentration for detecting the effect of CRS.

For the GFAP+ cell depletion studies, after habituating the animals for 48-hours to the sucrose solution (1%), the sucrose consumption was measured for the three 24 hrs periods following DT administration. This protocol allowed for a continuous monitoring of sucrose consumption, over the three day-period of administration of the DT and for detection of the onset of the behavioral deficits induced by GFAP+ cell depletion. The data was expressed as daily percent sucrose consumption since sucrose intake increases with time, in particular in control animals as they are exposed to sucrose. On day 4, following 16hrs fluid deprivation a 1-hour sucrose test was performed, and sucrose intake was measured in mL. On Day 8, we performed an additional 24-hour sucrose consumption monitoring to determine maintenance of the DT effect before euthanizing the animals. Water consumption was also measured to control for experimental bias associated with fluid drive

For the GFAP+ cell enhancement studies animals underwent 48-hours habituation to sucrose solution (2%) during the baseline week of testing. Following a 16-hour fluid deprivation, sucrose intake was measured for 1 hour and then mice were put back on water. In subsequent weeks, the animals were re-habited to sucrose for 24 hours and retested in the sucrose consumption test (16hr fluid deprivation + 1 hour sucrose intake). The same test is repeated later in the week with water instead of sucrose, as a control for general fluid intake. In weeks 3-5 where CNO (5mg/kg) was added to the either sucrose and water that the animal are drinking at the time. Both the sucrose and water consumption tests were performed at least 18 hours after their last CRS session (*7*).

*Elevated Plus Maze (EPM):* This maze is a plus shaped apparatus that is elevated 55cm off the ground. It has two open arms (27x5cm), and two enclosed arms (27x5x15cm). Mice are placed into the maze and allowed to explore for 10min in a dimly lit room (~20 lux). Time and number of entries in the open arms are recorded from above with a mounted camera and quantified using AnyMaze software. This test is used as a measure of anxiety-like behavior (*5*).

*Open Field (OF):* Mice are placed in an open novel arena (70x70x33cm) and are allowed to explore for 15mins. Time and number of entries in the center zone (40x40cm) are recorded from above with a mounted camera and quantified using AnyMaze software. This test is used as a measure of anxiety-like behavior (*5*).

*Novelty Supressed Feeding (NSF):* The NSF test begins after a 16-hour food deprivation period. The test then takes place in a novel arena (45×30×27cm) with a single food pellet placed in the centre of the cage under dimly lit conditions (lux ~20). The animal is then manually monitored and latency to feed on the pellet is recorded. The animals are then placed back in their home cage where a food pellet is placed in the centre and home cage latency to feed is recorded. The home cage test is performed to control for experimental bias associated with appetite drive. This test is used as an assessment of anxiety-like behavior (*5*).

*Novelty Induced Hypophagia (NIH):* The NIH test begins with three days of habituation to 30% sweetened milk. On the fourth day, a small dish with ~1mL of the sweetened milk is placed in the mouse home cage and homecage latency to drink the milk is recorded. The next day mice are placed in a novel home cage without any bedding and a small dish with ~1mL of the sweetened milk is placed in the cage and latency to drink the milk is then recorded. This test is a measure of anhedonia-like behavior (*4*).
*Forced Swim Test (FST):*

In this test mice are placed in a 5L glass beaker filled with 20cm high water and recorded from the side for 10 minutes. Time spent immobile was manually recorded from 2-10mins in 4-minute bins. In this test, time spent immobile is used to measure the animal's level of helplessness. This test is used as a measure of depressive-like behavior (*4*).

*Locomotor Activity (LM):* The locomotor activity test is used as a control for potential ambulatory bias. In this test, mice are again placed in a homecage-like arena and allowed to freely roam for 1-hour. Distance travelled in metres was quantified AnyMaze.

*PhenoTyper Test (PT):* We used the PhenoTyper test as a measure of conflict anxiety-like behavior in mice that has been shown to give consistent and reliable results over multiple weeks of testing and specific to mice subjected to stress (*5, 7, 9, 11*). The test utilizes the PhenoTyper (Noldus, Leesburg, VA, USA) apparatus, which consists of a home cage like arena (30x30cm) with a shelter, food, and water zone. The animals are placed into the apparatus weekly during their dark cycle period (7pm-7am) at least 1 hour after the second restraint session of the day. The conflict occurs when a light is automatically switched on at 11pm for 1 hour causing the mice to hide in their shelter. CRS mice were shown to display anxiety -like behavior, by continuing to hide in the shelter and avoid the remaining areas of the arena even after the light turns off (*5*). This residual avoidance is calculated for each animal using an equation as defined by Prevot et al., (*5*). Since food zone data usually mirrors data obtained in the shelter zone, only data based on time spent in the shelter zone were used in this study.

*Coat State Assessment (CS):* We performed weekly coat state assessments based on the protocol by Yalcin et al. (*12*), before the CRS sessions of the day. In short, the coat of 7 body parts (i.e., the head, neck, dorsal coat, ventral coat, tail, forepaws, and hind paws) were given a score of 0, 0.5, or 1 from well-kept to unkempt. The scores were then summated to determine the coat state degradation score of the animal's coat state.

*Anhedonia and Anxiety Z-Score:* To summarize our findings, we used an integrative z-scoring method to compile the anhedonia- or anxiety-like readouts into 1 score for each behavioral dimension per animal. The z-score is calculated first for each individual test using a function of the standard deviation and average of the control group for each sex. For the GFAP+ cell depletion study, anxiety z-score encompassed average z-score performances measured in the EPM, OF, and NSF. For the anhedonia z-score average z-score performances in day 8 sucrose consumption test and NIH were used. For the GFAP+ cell activity enhancement study anxiety z-score encompassed average z-score performances measured on the last week of CRS in the RA in the PhenoTyper and NSF. Last week sucrose consumption and NIH z-score were averages to calculate the anhedonia z-score. We used this approach in several studies assessing CRS and UCMS effects on anhedonia- and anxiety-like behaviors separately (*4, 13*).

*Immunochemistry:*

*Immunocytochemistry:* Primary astroglia cultures from wells containing coverslips were fixed using 4% paraformaldehyde in phosphate buffered saline (1XPBS) for 1-hour and washed with 1XPBS TritonX (0.3% PBS-T). Cells were then incubated for 30 mins in normal goat serum (10% NGS) in PBS-T, then with primary antibodies against GFP and Flag diluted in 3% NGS in PBS-T for 24 hours (4°C). Cells were then washed with 1XPBS and incubated in fluorescent secondary antibodies for 2 hours at room temperature and then washed again. Coverslips were then mounted on slides with Vectashield anti-fade mounting media (Vector labs) containing DAPI and visualized using confocal microscopy. References and concentrations of the primary and second antibodies can be found in Table.S1.

*Immunohistochemistry*: For immunohistochemistry, animals were perfused under anesthesia (Avertin, 125 mg/kg, i.p.) using 1XPBS (~30ml) and then 4% paraformaldehyde in 1XPBS (~100ml) via a Pharmacia pump (Cole-Parmer, Montreal, Canada). Brains were then dissected and post-fixed in 4% paraformaldehyde (overnight at 4°C), then cryoprotected in a 30% sucrose 1XPBS solution (48 hours at 4°C). Subsequently, the brains were frozen on dry ice before being cryosectioned using a cryostat (Leica CM1950, Buffalo Grove, IL, USA). Sections were 40-μm-thick containing the PFC (AP 2.3–1.4 mm from Bregma) and then placed in cryoprotectant (30% sucrose, 1% polyvinylpyrrolidone-40, 30% ethylene glycol) for storage at −20°C until immunochemistry.

The free-floating sections were first washed in 1XPBS, and then in TritonX (0.3-0.5% PBS-T) in 1XPBS. Sections were then incubated (1 hour or 30 mins) in serum (20% NGS or 10% normal donkey serum (NDS)) in PBS-T, then with primary antibodies (diluted in 3-6% NGS in

PBS-T or 3% NDS in PBS-T) for 24 hours at 4°C. After several washes with 1XPBS, sections incubated in the secondary antibodies for 2 hours at room temperature and then washed again in 1XPBS. Sections were then mounted on slides and cover slipped using with Vectashield anti-fade mounting media (Vector labs). The immunochemistry was done to validate the specificity of GFAP+ cell depletion virus (GFP and GFAP), extent of the depletion (GFP and Flag), selectivity of GFAP+ cell activation virus (GFAP, mCherry, and NeuN), selectivity of GFAP+ fiberphotometry virus (GFAP, GFP for GCaMP, and NeuN), and validation of chronic activation of GFAP+ cell (Fosb, GFAP, and mCherry). The concentration and combinations of antibodies employed can be found in Table.S1.

*Imaging and Quantification:*

For the GFAP+ cell depletion study, images of the immunostaining were taken using a Zeiss Axiovert LSM510 confocal microscope for one section per animal at the coordinates covering the site of viral infusion (AP+ 2-2.5). Cellular density of Flag+ cells was quantified at 20X, 5 z stacks per image, 0.1 μm z-step size. The number of Flag+ cells was quantified using Fiji Image J within a 0.5x0.5mm square and expressed in number of cells per $mm^2$.

For the GFAP+ cell activity enhancement study, images were captured using a Olympus IX83 inverted microscope equipped with a spinning disk confocal unit and Hamamatsu Orca-Flash4.0 V2 digital CMOS camera at 20X magnification in tandem with SlideBook 6 imaging software (Intelligent Imaging Innovations, Denver, CO). Two images per animal were taken from sections of the infected regions of PFC (40 z stacks per image, 0.1 μm z-step size). All images were taken at the same exposure for quantification (*6*). Fosb intensity of infected astroglia was quantified using the Fiji ImageJ fluorescence intensity software (*14*). Using the mCherry and GFAP channels, 10 GFAP+/mCherry+ astroglia were hand-traced without reference to the Fosb channel. Intensity of Fosb (raw integrated density) was quantified within those cells, then normalized for traced area of selected and the mean fluorescence of a background region. Total corrected cellular fluorescence (TCCF) was calculated as TCCF = raw integrated density – (area of selected cell × mean fluorescence of background) (*15*).

All quantifications were performed by an experimentation blinded to the slide code.

### SUPPLEMENTARY RESULTS

*In Vitro Cell Density:*

We confirmed the specificity of the designed viral construct *in vitro* in primary astrocyte cultures. Transfected cells expressed green fluorescence protein (GFP) when astroglia were generated from GFAPcre- pups and diphtheria toxin receptor fused with flag (DTRflag) was only present in cultures from GFAPcre+ pups (Supplementary Figure 1). Following 1 hour application of diphtheria toxin (DT), we found a 35% reduction of cell survival. Specifically, there was a significant effect of genotype ($F_{(1,20)}$=26.2,p<0.0001), treatment ($F_{(1,20)}$=24.6,p<0.0001), and genotype*treatment ($F_{(1,20)}$=19.9,p<0.001), explained by a reduction of cell density in with cre+ astroglia treated with DT compared to cre+ and cre- astroglia treated with PBS (p<0.0001)(Fig.S1d).

*Supplementary data collected for the GFAP+ cell depletion studies*:

Water consumption, home cage latency to feed in the NSF, home cage latency to drink in the NIH, distance travelled in the LM test, for each DT experiment can be found in Table.S2.

*Supplementary data collected during weekly behavioral testing of the GFAP+ cell activity enhancement study:*

*Baseline:* At baseline, we verified that mouse groups showed no difference in sucrose consumption ($t_{(1,42)}$=0.1; Fig.S5a), coat state degradation ($t_{(1,42)}$=0.3; Fig.S5b), or RA ($t_{(1,42)}$=0.3; Fig.S5d) compared to no CRS controls. For hourly recordings of time in the shelter zone, we found a significant main effect of time ($F_{(1,42)}$ = 51.8, p<0.0001), but no stress ($F_{(1,42)}$ = 0.1) or stress* time interaction ($F_{(1,12)}$ = 0.2)(Fig.S5c). Analysis of sex effects can be found in Table S3.

*Week 1:* After 1 week of stress, mice showed a significant effect of stress in sucrose consumption ($t_{(1,42)}$=8.7, p<0.05; Fig.S5e), coat state degradation ($t_{(1,42)}$=17.5, p<0.0001; Fig.S5f), and RA ($t_{(1,42)}$=20.3, p<0.0001; Fig.S5h) compared to no CRS controls. For hourly recordings of time in the shelter zone, there was a significant effect of time ($F_{(1,42)}$ = 67.0, p<0.0001), stress ($F_{(1,42)}$ = 15.0, p<0.001), and a stress* time interaction ($F_{(1,12)}$ = 3.6, p<0.0001)(Fig.S5g). Analysis of sex effects can be found in Table S3.

*Week 2:* Data can be found in the main text and analysis of sex effects can be found in Table S3.

*Week 3:* There was no significant effect of stress ($F_{(1,40)}$ = 0.1), drug ($F_{(1,40)}$ = 3.2), or stress*drug interaction ($F_{(1,40)}$ = 0.9) on sucrose consumption (Fig.S5i). In both coat state degradation and RA, we found a significant main of stress (CS:$F_{(1,40)}$ = 11.5, p <0.01; RA:$F_{(1,40)}$ = 32.7, p <0.0001), but no effect of drug (CS: $F_{(1,40)}$ = 0.3; RA: $F_{(1,40)}$ = 0.6), or drug*stress interaction (CS: $F_{(1,40)}$ = 0.5, p>0.05; RA: $F_{(1,40)}$ = 0.4, p>0.05), explained by a significant difference between homecage controls and CRS+No CNO (CS: p<0.01; RA: p<0.0001) and CRS+CNO (CS: p=0.06; RA: p<0.001)(Fig.S5j,l). Analysis of time spent in the shelter zone, revealed a significant main effect of stress ($F_{(1,40)}$ = 32.9, p <0.0001), time ($F_{(1,40)}$ = 45.1, p<0.0001), and stress*time interaction($F_{(1,12)}$ = 6.5, p<0.0001), but no effect of drug ($F_{(1,40)}$ = 0.1), stress*drug interaction ($F_{(1,12)}$ = 2.6), drug*time interaction ($F_{(1,12)}$ = 1.1), or stress*drug*time interaction ($F_{(1,12)}$=0.8)(Fig.S5k). Analysis of sex effects can be found in Table S3.

*Week 4:* Analysis of sucrose consumption, revealed no effect of stress ($F_{(1,40)}$ = 4.9, p<0.05), drug ($F_{(1,40)}$ = 0.8, p>0.05), or stress*drug interaction ($F_{(1,40)}$ = 0.1, p>0.05)(Fig.S5m). In addition, when looking at coat state degradation and RA there was a significant main effect of stress (CS: $F_{(1,40)}$ = 21.1, p <0.0001; RA: $F_{(1,40)}$ = 85.6, p <0.0001), but no drug (CS: $F_{(1,40)}$ = 0.1; RA: $F_{(1,40)}$ = 0.2), or drug*stress interaction (CS: $F_{(1,40)}$ = 0.5; RA: $F_{(1,40)}$ = 0.1). This is explained by a significant increase in coat state degradation and RA between homecage controls and No CRS+NoCNO (CS & RA: p<0.0001) and CRS+CNO (CS: p<0.01; RA: p<0.0001) (Fig.S5n,p). Analysis of time spent in the shelter zone revealed a significant main effect of stress ($F_{(1,40)}$ = 75.8, p <0.0001), time ($F_{(1,40)}$ = 63.8, p<0.0001), stress*time interaction ($F_{(1,12)}$ = 10.1, p<0.0001), and stress*drug*time interaction ($F_{(1,12)}$=2.0; p<0.05), but no effect of drug ($F_{(1,40)}$ = 0.2), stress*drug interaction ($F_{(1,12)}$ = 0.5), or drug*time interaction ($F_{(1,12)}$ = 0.9)(Fig.S5o). Analysis of sex effects can be found is Table S3.

*Week 5:* Data can be found in the main text and analysis of sex effects can be found in Table S3.

Water consumption, home cage latency to feed in the NSF, home cage latency to drink in the NIH, distance travelled in the LM test, for the GFAP+ cell activity enhancement experiment can be found in Table.S2.

*Supplementary data collected during weekly behavioral testing of the CNO control study in CRS animals:*

*Baseline:* Throughout this experiment, for simplicity sucrose consumption data are expressed in percent sucrose consumption calculated as sucrose intake over total fluid (sucrose+water) intake. Analysis of percent sucrose consumption (SC), CS and RA revealed no effect of stress (SC: $F_{(1,36)}$ = 0.1; CS: $F_{(1,36)}$ = 0.9; RA: $F_{(1,36)}$ = 0.1), drug (SC: $F_{(1,36)}$ = 0.8; CS: $F_{(1,36)}$ = 0.9; RA: $F_{(1,36)}$ = 0.1), or stress*drug interaction (SC: $F_{(1,36)}$ = 0.6; CS: $F_{(1,36)}$ = 0.1; RA: $F_{(1,36)}$ = 0.1). Analysis of time spent in the shelter zone also revealed no effect of stress ($F_{(1,36)}$ = 0.1), drug ($F_{(1,36)}$ = 0.3), stress*drug interaction ($F_{(1,36)}$ = 0.1), but a significant effect of time ($F_{(1,12)}$ = 31.1, p<0.0001). There was also no time*stress ($F_{(1.12)}$=0.8), time*drug ($F_{(1.12)}$=0.6), or stress*drug*time interaction ($F_{(1.12)}$=1.0).

*Week 1:* Analysis of percent sucrose consumption test revealed no effect of stress ($F_{(1,36)}$ = 0.1), drug ($F_{(1,36)}$ = 0.6), or stress*drug interaction ($F_{(1,36)}$ = 0.4). In the CS test, we found a significant effect of stress ($F_{(1,36)}$ = 29.0, p<0.0001) but no drug ($F_{(1,36)}$ = 0.2), or drug*stress interaction ($F_{(1,36)}$ = 0.1) explained by a significant increase in coat state degradation between homecage controls and CRS+No CNO as well as CRS+CNO animal groups (p<0.001). Analysis of RA showed a significant main effect of stress ($F_{(1,36)}$ = 37.3, p>0.0001), drug ($F_{(1,36)}$ = 4.3), but no stress*drug interaction ($F_{(1,36)}$ = 1.1), explained by an increase in RA between No CRS+CNO and CRS+CNO (p<0.001), NoCRS+No CNO and CRS+No CNO groups (p<0.0001), and a decrease in RA between CRS+No CNO and CRS+CNO groups (p<0.05). On time spent in the shelter zone there was again a significant effect of stress ($F_{(1,36)}$ = 40.1, p<0.0001), no drug ($F_{(1,36)}$ = 1.1), stress*drug interaction ($F_{(1,36)}$ = 0.1). There was a significant effect of time ($F_{(1,12)}$ = 44.8, p<0.0001), time*stress ($F_{(1.12)}$=2.9,p<0.001), but no time*drug ($F_{(1.12)}$=1.6), or stress*drug*time interaction ($F_{(1.12)}$=1.1), explained by both CRS groups spending more time in the shelter following the light challenge as compared to their respective homecage groups (p<0.0001).

*Week 2:* Analysis of percent sucrose consumption on week 2 revealed no effect of stress ($F_{(1,36)}$ = 0.7), drug ($F_{(1,36)}$ = 1.8), or stress*drug interaction ($F_{(1,36)}$ = 0.1). When looking at CS and RA, we found a significant main effect of stress (CS: $F_{(1,36)}$ = 28.2, p<0.0001; RA: $F_{(1,36)}$ = 57.1, p<0.0001) but no effect of drug (CS: $F_{(1,36)}$ = 1.5, p>0.05; RA: $F_{(1,36)}$ = 0.4, p>0.05) or stress*drug interaction (CS: $F_{(1,36)}$ = 1.0, p>0.05; RA: $F_{(1,36)}$ = 0.6, p>0.05) explained by a significant increase in coat state degradation and RA between homecage groups and CRS+CNO (CS & RA: $p<0.0001$) as well as CRS+No CNO (CS: $p<0.01$; RA: $p<0.0001$). For time spent in the shelter zone there was a significant effect of stress ($F_{(1,36)} = 67.3$, $p<0.0001$), but no drug ($F_{(1,36)} = 0.1$), or stress*drug interaction ($F_{(1,36)} = 0.8$), although there was a significant effect of time ($F_{(1,12)} = 46.2$, $p<0.0001$). There was also a time*stress ($F_{(1.12)}=5.2$,$p<0.0001$), but no time*drug ($F_{(1.12)}=1.1$), or stress*drug*time interaction ($F_{(1.12)}=0.6$), explained by CRS groups spending more time in the shelter following the light challenge ($p<0.0001$).

*Week 3:* Analysis of percent sucrose consumption in week 3 showed an effect of stress ($F_{(1,36)} = 5.0$, $p<0.05$), drug ($F_{(1,36)} = 4.0$, $p=0.05$), and stress*drug interaction ($F_{(1,36)} = 9.5$, $p<0.05$) explained by a significant decrease in percent sucrose consumption between NoCRS+CNO and CRS+CNO ($p<0.001$) and between CRS+No CNO and CRS+CNO ($p<0.01$). Further analysis of CS and RA revealed a significant main effect of stress (CS: $F_{(1,36)} = 56.3$, $p<0.0001$; RA: $F_{(1,36)} = 18.7$, $p<0.0001$) but no effect of drug (CS: $F_{(1,36)} = 0.1$; RA: $F_{(1,36)} = 0.4$) or stress*drug interaction (CS: $F_{(1,36)} = 0.1$; RA: $F_{(1,36)} = 0.7$). This is explained as a significant increase in coat state degradation and RA between homecage controls and CRS+No CNO (CS & RA: $p<0.001$) and CRS+CNO (CS: $p<0.0001$; RA: $p<0.05$). Hourly time spent in the shelter zone analysis revealed a significant effect of stress ($F_{(1,36)} = 28.4$, $p<0.0001$) and time ($F_{(1,12)} = 30.6$, $p<0.0001$), but no drug ($F_{(1,36)} = 0.3$), or stress*drug interaction ($F_{(1,36)} = 1.2$). There was also a time*stress ($F_{(1.12)}=3.0$,$p<0.001$), but no time*drug ($F_{(1.12)}=1.2$), or stress*drug*time interaction ($F_{(1.12)}=1.1$), explained by CRS groups spending more time in the shelter following the light challenge ($p<0.0001$).

*Week 4:* There was no effect of stress ($F_{(1,36)} = 1.0$), drug ($F_{(1,36)} = 0.3$), or stress*drug interaction ($F_{(1,36)} = 0.3$) on percent sucrose consumption. Analysis of CS and RA showed an effect of stress (CS: $F_{(1,36)} = 56.8$, $p<0.0001$; RA: $F_{(1,35)} = 15.2$, $p<0.001$), but no effect of drug (CS: $F_{(1,36)} = 0.9$; RA: $F_{(1,35)} = 0.3$) or stress*drug interaction (CS: $F_{(1,36)} = 0.1$; RA: $F_{(1,35)} = 0.5$). This is explained by a significant increase in coat state degradation and RA between homecage groups and CRS+No CNO (CS: $p<0.01$; RA: $p<0.0001$) and CRS+CNO (CS: $p<0.05$; RA: $p<0.0001$). For time spent in the shelter zone there was a significant effect of stress ($F_{(1,35)} = 35.6$, $p<0.0001$), but no drug ($F_{(1,35)} = 0.1$), or stress*drug interaction ($F_{(1,35)} = 3.6$), although there was a significant effect of time ($F_{(1,12)} = 34.7$, $p<0.0001$). There was also a time*stress ($F_{(1.12)}=3.8$, $p<0.0001$), but no time*drug ($F_{(1.12)}=0.7$), or stress*drug*time interaction ($F_{(1.12)}=0.6$), explained by CRS groups spending more time in the shelter following the light challenge ($p<0.0001$). Due to technical complications with the PhenoTyper, data from one mouse in the CRS+CNO group were lost on this week.

*Week 5:* There is no effect of stress ($F_{(1,36)} = 0.4$), drug ($F_{(1,36)} = 0.3$), or stress*drug interaction ($F_{(1,36)} = 0.4$) on percent sucrose consumption. Analysis of CS and RA showed an effect of stress (CS: $F_{(1,36)} = 39.8$, $p<0.0001$; RA: $F_{(1,36)} = 17.0$, $p<0.001$) but no effect of drug (CS: $F_{(1,36)} = 2.2$; RA: $F_{(1,36)} = 0.8$) or stress*drug interaction (CS: $F_{(1,36)} = 1.7$; RA: $F_{(1,36)} = 2.7$). This is explained by a significant increase in coat state degradation and RA between homecage groups and CRS+No CNO (CS: $p<0.01$; RA: $p<0.001$) and CRS+CNO (CS: $p<0.0001$; RA: $p=0.09$). Additionally, the CS analysis revealed a significant difference between CRS+No CNO and CRS+CNO ($p=0.05$) such that mice exposed to CNO show a greater amount of CS degradation. For time spent in the shelter zone there was a significant effect of stress ($F_{(1,36)} = 27.3$, $p<0.0001$), but no drug ($F_{(1,36)} = 1.2$, $p>0.05$), or stress*drug interaction ($F_{(1,36)} = 0.2$, $p>0.05$), although there was a significant effect of time ($F_{(1,12)} = 50.5$, $p<0.0001$). There was also a time*stress ($F_{(1.12)}=3.4$,$p<0.0001$), but no time*drug ($F_{(1.12)}=0.9$), or stress*drug*time interaction ($F_{(1.12)}=1.9$), explained by CRS groups spending more time in the shelter following the light challenge ($p<0.0001$).

Repeated-measures analysis over the full 5 week period on percent sucrose consumption, revealed no significant effect of stress ($F_{(1,36)} = 0.2$), drug ($F_{(1,36)} = 1.7$), or stress*drug interaction ($F_{(1,36)} = 0.1$), a significant effect of time ($F_{(5,36)} = 5.2$, $p<0.001$), but no time*stress ($F_{(5,36)} = 0.2$) or time*drug interaction ($F_{(5,36)} = 0.7$) (Fig.S6a). Data from collected in animal subjected to CRS and CNO for 3 weeks on CS (Fig.S6b), hourly time spent in the shelter zone (Fig.S6c), and RA (Fig.S6d) are highlighted as it is the most relevant time point in comparison to the GFAP+ cell activity enhancement studies. Throughout the experiment there was little to no effect of drug or drug*stress interactions throughout the entire study and that there were no sex*drug interactions on any of the behavior at each time point (data not shown). Main effect of sex and sex*stress interactions mirrored the findings obtained for the GFAP+ cell activity enhancement study. No difference between groups were found in LM activity throughout the experiment.

### SUPPLEMENTARY FIGURE LEGENDS

**Fig. S1 – Conditional expression of diphtheria toxin receptor (DTR) and cell ablation in primary astrocyte culture from glial fibillary acidic protein (GFAP) cre+ and cre- mice**. (a) Astrocytes (blue-DAPI) were generated from GFAPcre- mice, maintained *in vitro*, and transfected with GFP-DIOCMV-DTRflag plasmid. The CMV promoter drives the expression of green fluorescent protein (GFP) in transfected cells (white arrows). In these conditions no DTRflag is expressed. (b-c) Astrocytes (blue-DAPI) were generated from GFAPcre+ mice, maintained *in vitro*, and transfected with GFP-DIOCMV-DTRflag plasmid. The CMV promoter drives the expression of the DTRflag in transfected cells (empty arrows). GFP immunostaining showed that some cells express very low levels of GFP (white arrow), most infected cells express only DTRflag (empty arrow) (a-c) Scale=10um. (d) MTT survival assay was used to assess cell viability in GFAPcre+ and GFAPcre- cell cultures transfected with GFP-DIOCMV-DTRflag plasmid and treated or not with DT for 1 hour. Data are presented as percent cell survival and mean ± SEM. ****$p < .0001$ as compared to PBS.

**Fig.S2: Effects of cortical infusion of AAV5-GFP-DIOCMV-DTRflag and i.p. injections of diphtheria toxin (DT) in GFAPcre- mice.** (a-f) GFAPcre- mice were infused with AAV5-GFP-DIOCMV-DTRflag in the prefrontal cortex. Following three weeks recovery, mice were administered with several doses of DT (circle - 0, square - 0.1, triangle - 5, and inverted triangle - 20µg/kg). Percent sucrose consumption in the sucrose (ST) was measured on day 1 (a), day 2 (b), and day 3 (c) for 24 hours. Mice were also tested in the novelty suppressed feeding (NSF)(d), novelty induced hypophagia (NIH )(e), and forced swim test (FST)(f). Additional behavioral assessments (LM), water intake, and home-cage latency to feed and drink) can be found in Supplementary Table 2. Data are presented as individual animals and mean ± SEM.

**Fig.S3: Effects of cortical infusion of AAV5-GFP-mDIOCMV-DTRflag and i.p. injections of diphtheria toxin (DT) in GFAPcre+ mice.** (a-f) GFAPcre+ mice were infused with AAV5-GFP-mDIOCMV-DTRflag in the prefrontal cortex. Following three weeks recovery, mice were administered with of saline (circle - 0) or DT( inverted triangle - 20µg/kg). Percent sucrose consumption in the sucrose (ST) was measured on day 1 (a), day 2 (b), and day 3 (c) for 24 hours. Mice were also tested in the novelty suppressed feeding (NSF)(d), novelty induced hypophagia (NIH )(e), and forced swim test (FST)(f). Additional behavioral assessments (LM), water intake, and home-cage latency to feed and drink) can be found in Supplementary Table 2. Data are presented as individual animals and mean ± SEM.

**Fig.S4: Effects of striatal infusion of AAV5-GFP-DIOCMV-DTRflag and i.p. injections of diphtheria toxin (DT) in GFAPcre+ and cre- mice.** (a-e) GFAPcre+ and wildtype (WT) littermates were infused with AAV5-GFP-DIOCMV-DTRflag in the striatum. Following three weeks recovery, mice were administered with of saline (circle - 0) or DT( inverted triangle - 20µg/kg). Percent sucrose consumption in the sucrose (ST) was measured on day 1 (a), day 2 (b), and day 3 (c) for 24 hours. Mice were also tested in the novelty suppressed feeding (NSF)(d), and novelty induced hypophagia (NIH )(e). Additional behavioral assessments (LM), water intake, and home-cage latency to feed and drink) can be found in Supplementary Table 2. Data are presented as individual animals and mean ± SEM.

**Fig.S5: Cortical Glial Fibrillary Acidic Protein (GFAP)+ cell activity enhancement reverses chronic restraint stress (CRS)-induced anhedonia- but not anxiety-like behavior.** Mice infused in the PFC with the AAV5-GFAP-hM3D(Gq)-mCherry virus and tested for baseline behavior (a-d). Mice were then subjected to CRS or not for 2 weeks (week 1 data: e-h, week 2 data: see figure 3b-e) and treated with clozapine-n-oxide (CNO) or not for the remaining 3 weeks (week 3:i-l, week 4:m-p, week 5:see figure 3f-i). Mice were tested weekly for sucrose consumption in the sucrose test (ST) (a,e,i,m), and coat state degradation in coat state (CS) (b,f,j,n). Hourly time spent in shelter (c,g,k,o - *p< .05, **p< .01, ***p<.001, and ****p<.0001: No CRS+No CNO vs. CRS+No CNO ; #p< .05, ##p< .01, ###p<.001,and ####p<.0001: No CRS+CNO vs. CRS+CNO) and RA (d,h,l,p) in the PhenoTyper Test (PT) were also monitored weekly. Data are presented as mean ± SEM. Between group significant differences are marked as *p< .05, **p< .01, ***p<.001, and ****p<.0001: No CRS+No CNO vs. CRS+No CNO; #p< .05, ##p< .01, ###p<.001,and ####p<.0001: No CRS+CNO vs. CRS+CNO

**Fig.S6: Chronic restraint stress (CRS) and chronic clozapine-N-oxide (CNO) administration in C57Bl/6.** (a) Percent sucrose consumption of WT mice subjected or not to 5 weeks of CRS and treated or not with CNO were measured at baseline and weekly during CRS exposure. (b-d) The effects of administration of CNO for 3 weeks were assessed on coat state (b), in the PhenoTyper Test on the hourly time spent in the shelter zone (c - *p< .05, **p< .01, and ****p<.0001: No CRS+No CNO vs. CRS+No CNO ; #p<0.05, ##p< .01: No CRS+CNO vs. CRS+CNO, #p<0.05, &&p< .01: CRS+NoCNO vs. CRS+CNO) and residual avoidance (RA)(d). Data are presented as mean ± SEM. Between group significant differences are marked as *p< .05, **p< .01, ***p<.001, and ****p<.0001: No CRS+No CNO vs. CRS+No CNO; #p< .05, ##p< .01, ###p<.001,and ####p<.0001

**Fig.S7: Increased Fosb intensity in infected Cortical Glial Fibrillary Acidic Protein (GFAP)+ cells following CNO administration.** Immunohistochemistry identifying GFAP+ cells (green), mCherry+ (red), and Fosb +(blue). White arrow indicates GFAP+/mCherry+/Fosb+ cells in No CNO conditions (a) and in an animal treated with CNO (b). Scale = 10um
