## Supplementary material for "Prefrontal Cortex Astroglia Modulate Anhedonia-like Behavior": Fig.S1, Fig.S2, Fig.S3, Fig.S4, Fig.S5, Fig.S6, Fig.S7, Table.S1, Table.S2, Table.S3 will link to this file

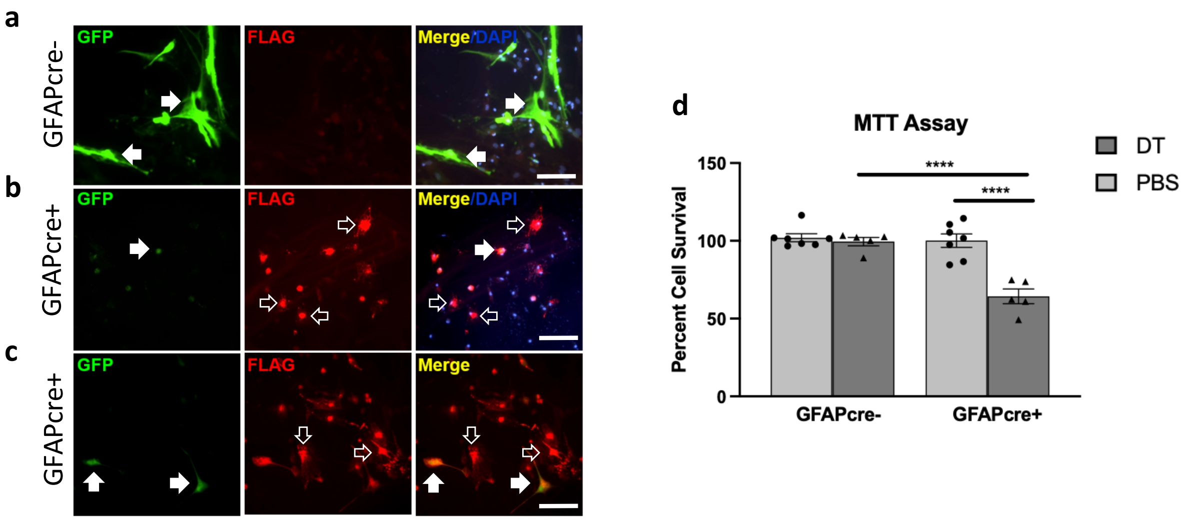
a
b
c
GFAPcre-
GFAPcre+
GFAPcre+
GFP
FLAG
Merge/DAPI
Merge
d
MTT Assay
DT
PBS
Percent Cell Survival
150
100
50
0
****
****
GFAPcre-
GFAPcre+

Fig.S2

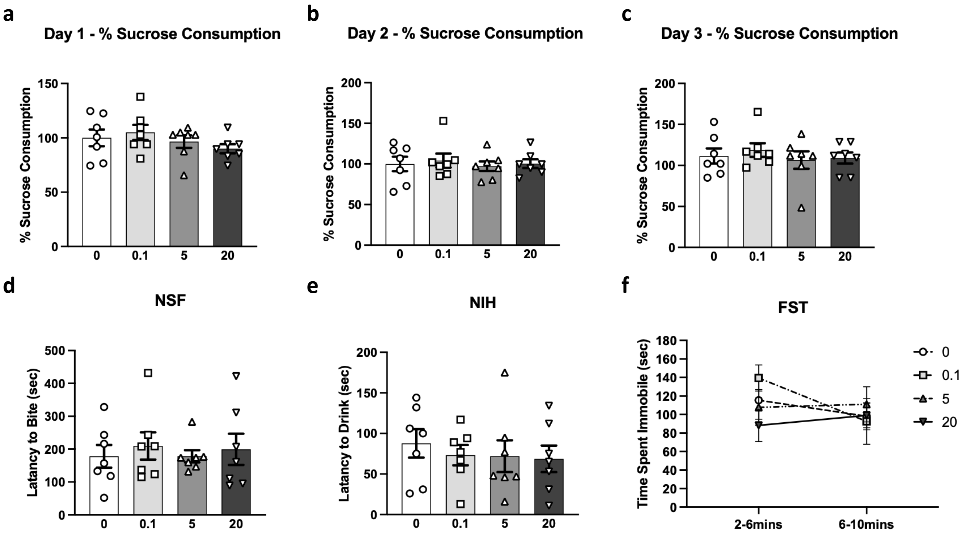

a
Day 1 - % Sucrose Consumption
% Sucrose Consumption
0
0.1
5
20
b
Day 2 - % Sucrose Consumption
% Sucrose Consumption
c
Day 3 - % Sucrose Consumption
% Sucrose Consumption
d
NSF
Latency to Bite (sec)
e
NIH
Latency to Drink (sec)
f
FST
Time Spent Immobile (sec)
2-6mins
6-10mins
0
0.1
5
20

Fig.S3

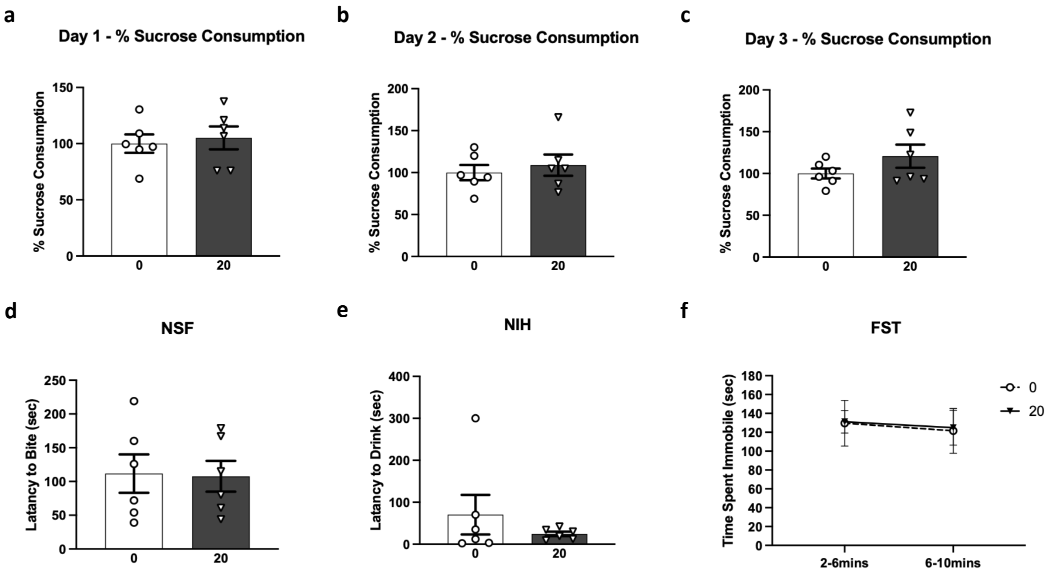

a
Day 1 - % Sucrose Consumption
% Sucrose Consumption
150
100
50
0
0
20
b
Day 2 - % Sucrose Consumption
% Sucrose Consumption
200
150
100
50
0
0
20
c
Day 3 - % Sucrose Consumption
% Sucrose Consumption
200
150
100
50
0
0
20
d
NSF
Latancy to Bite (sec)
250
200
150
100
50
0
0
20
e
NIH
Latancy to Drink (sec)
400
300
200
100
0
0
20
f
FST
Time Spent Immobile (sec)
180
160
140
120
100
80
60
40
20
0
2-6mins
6-10mins
0
20

Fig.S4

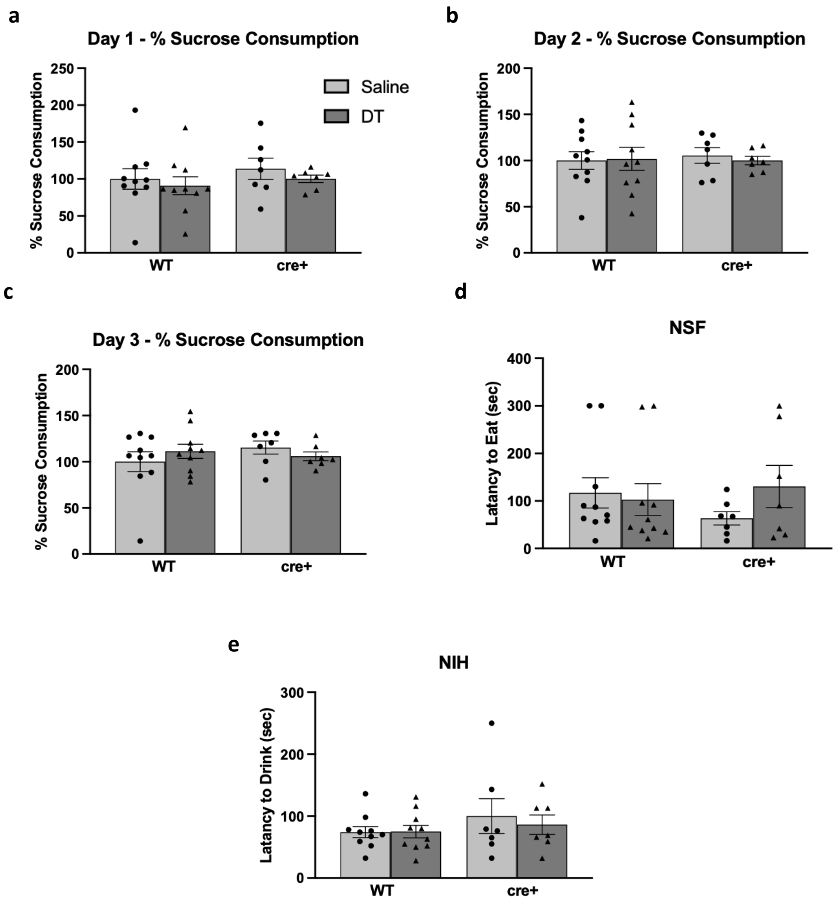
a
Day 1 - % Sucrose Consumption
% Sucrose Consumption
250
200
150
100
50
0
Saline
DT
WT
cre+
b
Day 2 - % Sucrose Consumption
% Sucrose Consumption
200
150
100
50
0
WT
cre+
c
Day 3 - % Sucrose Consumption
% Sucrose Consumption
200
150
100
50
0
WT
cre+
d
NSF
Latency to Eat (sec)
400
300
200
100
0
WT
cre+
e
NIH
Latency to Drink (sec)
300
200
100
0
WT
cre+

Fig.S5

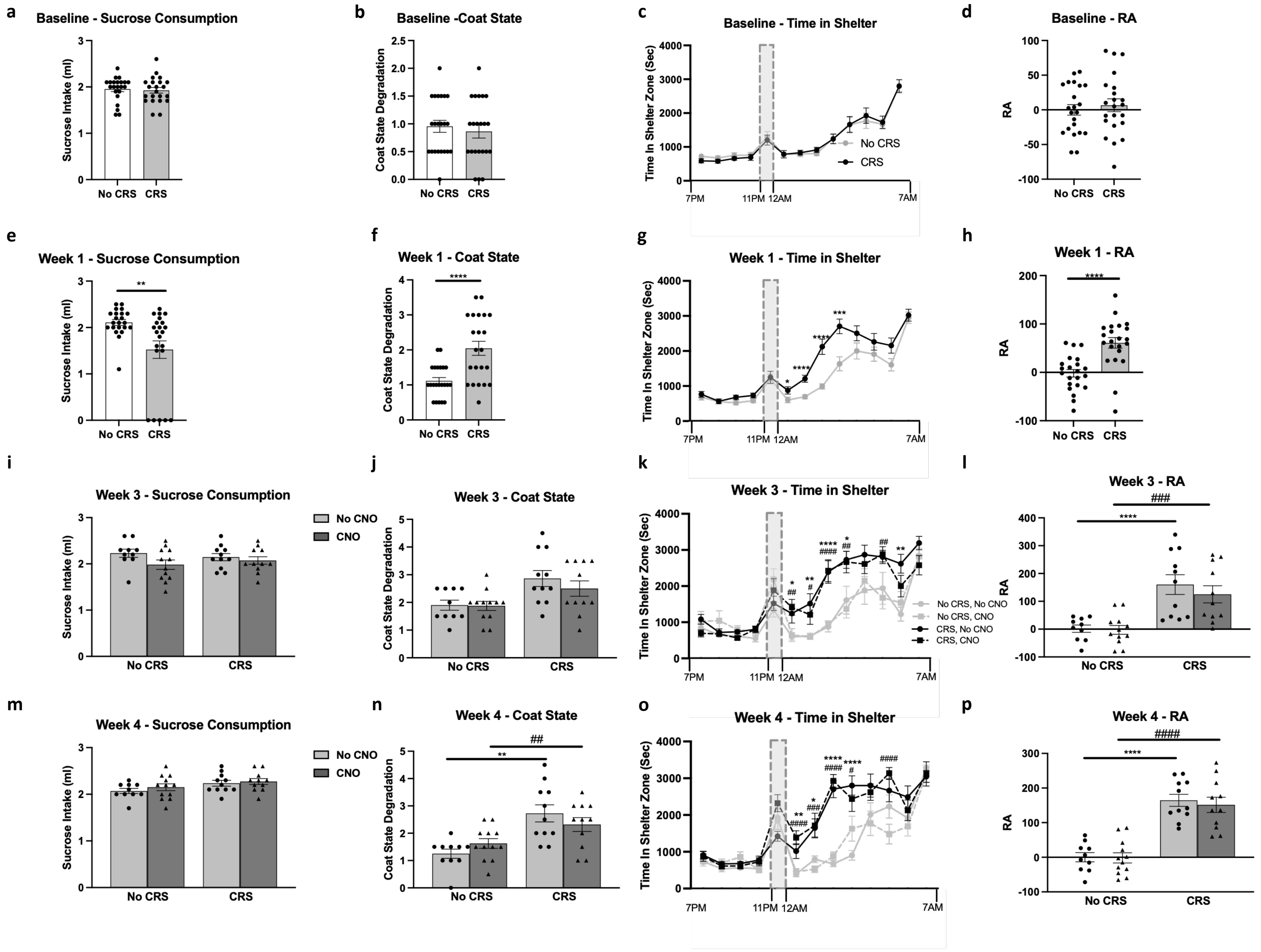
a
Baseline - Sucrose Consumption
Sucrose Intake (ml)
No CRS
CRS
b
Baseline -Coat State
Coat State Degradation
c
Baseline - Time in Shelter
Time In Shelter Zone (Sec)
No CRS
CRS
7PM
11PM
12AM
7AM
d
Baseline - RA
RA
e
Week 1 - Sucrose Consumption
f
Week 1 - Coat State
g
Week 1 - Time in Shelter
h
Week 1 - RA
i
Week 3 - Sucrose Consumption
No CNO
CNO
j
Week 3 - Coat State
k
Week 3 - Time in Shelter
No CRS, No CNO
No CRS, CNO
CRS, No CNO
CRS, CNO
l
Week 3 - RA
m
Week 4 - Sucrose Consumption
n
Week 4 - Coat State
o
Week 4 - Time in Shelter
p
Week 4 - RA

Fig.S6

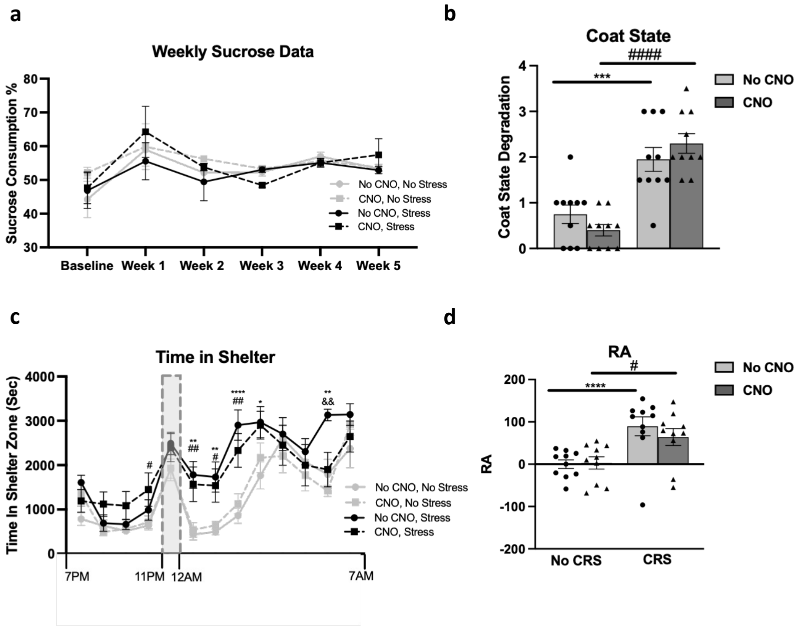
a
Weekly Sucrose Data
Sucrose Consumption %
80
70
60
50
40
30
Baseline
Week 1
Week 2
Week 3
Week 4
Week 5
No CNO, No Stress
CNO, No Stress
No CNO, Stress
CNO, Stress
b
Coat State
Coat State Degradation
####
***
No CNO
CNO
4
3
2
1
0
c
Time in Shelter
Time In Shelter Zone (Sec)
4000
3000
2000
1000
0
7PM
11PM
12AM
7AM
No CNO, No Stress
CNO, No Stress
No CNO, Stress
CNO, Stress
d
RA
#
****
No CNO
CNO
200
100
0
-100
-200
No CRS
CRS

Fig.S7

**a**

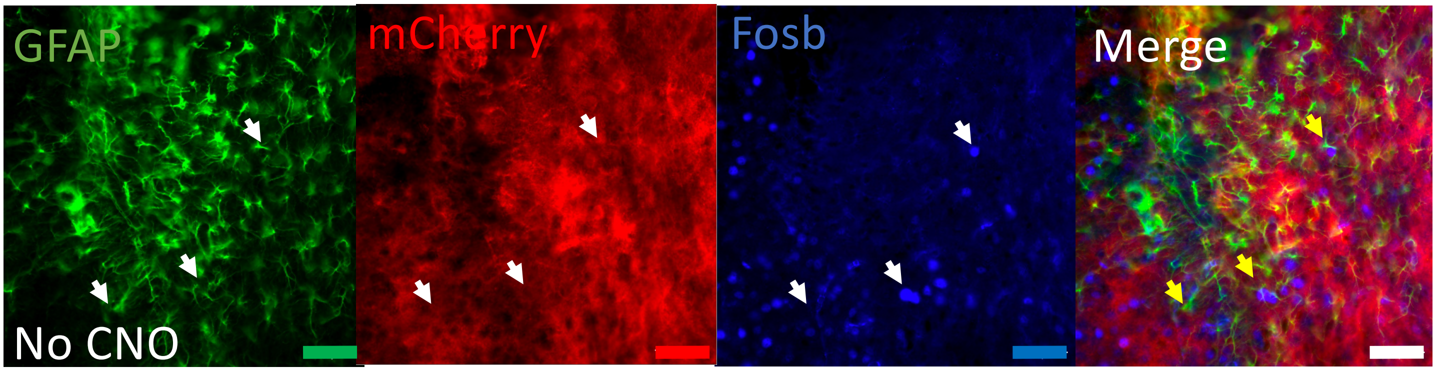


**b**

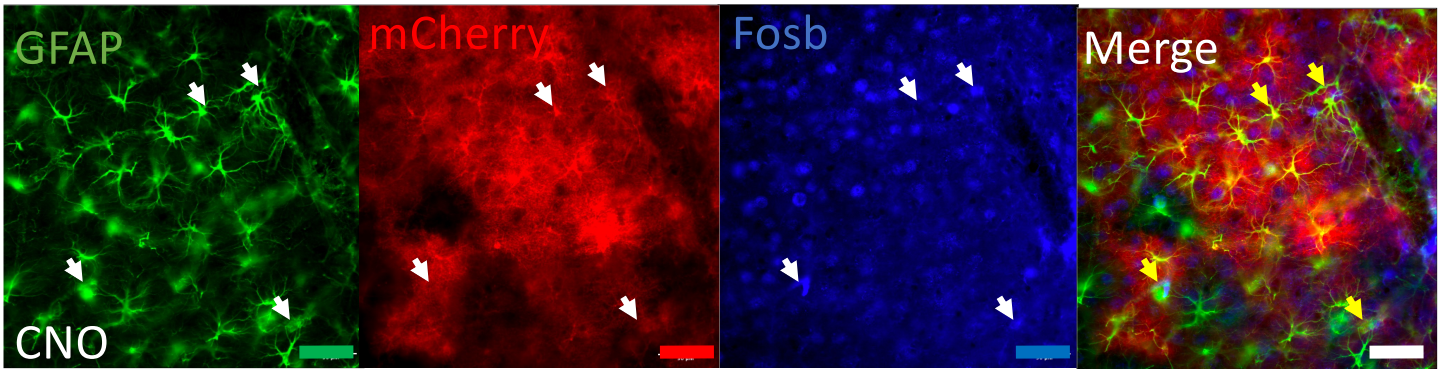

**Table 1: List of Antibodies**

| Purpose | Primary Antibody | Company/Catalog # | Secondary Antibody | Company/Catalog # |
|---|---|---|---|---|
| **Fosb Flouresence Intensity** | Chk α mCherry (1:1000) | Abcam – ab 205 402 | Goat α Chk AF 568 (1:500) | Invitrogen (thermofisher) ref A11041 |
| | Ms α FosB (1:100) | Abcam – ab 11959 | Goat α Ms AF 405 (1:100) | Invitrogen A31553 |
| | Rb α GFAP (1:100) | Dako - Z0334 | Goat α Rb AF 488 (1:250) | Invitrogen A11008 |
| **pAAV-GFAP-hM3D(Gq)-mCherry - Validation** | Chk α mCherry (1:1000) | Abcam – ab 205 402 | Goat α Chk AF 568 (1:250) | Invitrogen (thermofisher) ref A11041 |
| | Rb α GFAP (1:200) | Dako - Z0334 | Goat α Rb AF 488 (1:250) | Invitrogen A11008 |
| | Ms α NeuN (1:200) | Millipore - MAB377 | Goat α Ms AF 405 (1:250) | Invitrogen A31553 |
| **pZac2.1 gfaABC1D-lck-GCaMP6f virus** | Rb α GFAP (1:200) | Dako - Z0334 | Donkey α Rb AF 555 (1:200) | Invitrogen - A21572 |
| | Gt α GFP (1:200) | Rockland - 600-101-215 | Donkey α Gt AF 488 (1:200) | Invitrogen - A11055 |
| | GP α NeuN (1:200) | EMD Millipore - ABN90 | Donkey α GP AF 405 (1:200) | Biotium - 20376 |
| **AAV5-GFP-DIOCMV-DTRflag- In Vitro Validation** | Rb α Flag (1:500) | Sigma - F7425 | Goat α Rb AF 568 (1:500) | Thermofisher/Invitrogen - A11011 |
| | Chk α GFP (1:700) | Aves Lab - GFP1020 | Goat α Chk AF 488 (1:500) | Thermofisher/Invitrogen - A11039 |
| **AAV5-GFP-DIOCMV-DTRflag- In Vitro Validation** | Rb α Flag (1:250) | Sigma - F7425 | Goat α Rb AF 568 (1:200) | Thermofisher/Invitrogen - A11011 |
| | Chk α GFP (1:500) | Aves Lab - GFP1020 | Goat α Chk AF 488 (1:200) | Thermofisher/Invitrogen - A11039 |
| | Rb α GFAP (1:200) | Dako - Z0334 | Goat α Rb AF 568 (1:200) | Thermofisher/Invitrogen - A11011 |
| | Rb = Rabbit, Chk = Chicken, MS= Mouse, GT= Goat, Glial Fibillary Acidic Protein (GFAP), Green Fluorescent Protein (GFP) | | | |

**Table 2: Statistical analysis of water consumption, latency to eat and drink in homecage condition, and locomotor activity**

| Experiment | Control Test | Statistics (F,p) | Post-hoc analysis |
|---|---|---|---|
| **PFC Astroglial Abalation in GFAPcre+ Mice** | Day 5 Water Consumption | (F(3,22) =0.5, p>0.05) | N/A |
| | Latency to Drink in Homecage | (F(3,22) =1.2, p>0.05) | N/A |
| | Latency to Eat in Homecage | (F(3,22) =0.5, p>0.05) | N/A |
| | Locomotor | (F(3,22) =0.5, p>0.05) | N/A |
| **PFC Astroglial Abalation in GFAPcre+ Mice using mutant virus** | Day 5 Water Consumption | (F(1,10) =1.1, p>0.05) | N/A |
| | Latency to Drink in Homecage | (F(1,10) =0.9, p>0.05) | N/A |
| | Latency to Eat in Homecage | (F(1,10) =1.2, p>0.05) | N/A |
| | Locomotor | (F(1,10) =1.6, p>0.05) | N/A |
| **PFC Astroglial Abalation in WT Mice** | Day 5 Water Consumption | (F(3,24) =0.2, p>0.05) | N/A |
| | Latency to Drink in Homecage | (F(3,24) =1.7, p>0.05) | N/A |
| | Latency to Eat in Homecage | (F(3,24) =5.8, p<0.01)* | 5ug/kg vs. Vehicle *** |
| | Locomotor | (F(3,24) =2.2, p>0.05) | N/A |
| **Striatum Astroglial Abalation in GFAPcre+ Mice** | Day 5 Water Consumption | Genotype: (F(1,30) =0.5, p>0.05) | N/A |
| | | Treatment: : (F(1,30) =1.1, p>0.05) | N/A |
| | | Genotype*Treatment: (F(1,30) =0.1, p>0.05) | N/A |
| | Latency to Drink in Homecage | Genotype: (F(1,30) =0.3, p>0.05) | N/A |
| | | Treatment: : (F(1,30) =0.1, p>0.05) | N/A |
| | | Genotype*Treatment: (F(1,30) =0.1, p>0.05) | N/A |
| | Latency to Eat in Homecage | Genotype: (F(1,30) =0.1, p>0.05) | N/A |
| | | Treatment: : (F(1,30) =0.2, p>0.05) | N/A |
| | | Genotype*Treatment: (F(1,30) =3.8, p>0.05) | N/A |
| | Locomotor | Genotype: (F(1,30) =1.2, p>0.05) | N/A |
| | | Treatment: : (F(1,30) =0.3, p>0.05) | N/A |
| | | Genotype*Treatment: (F(1,30) =0.8, p>0.05) | N/A |
| **PFC Astroglial Enhancement** | Water Intake Baseline | (F(1,42) =0.4, p>0.05) | N/A |
| | Water Intake Week 1 | (F(1,42) =1.9, p>0.05) | N/A |
| | **Water Intake Week 2** | (F(1,42) =1.6, p>0.05) | N/A |
| | Water Intake Week 3 | Stress: (F(1,42) =2.3, p>0.05) | N/A |
| | | Drug: (F(1,42) =1.0, p>0.05) | N/A |
| | | Stress*Drug: (F(1,42) =0.1, p>0.05) | N/A |
| | Water Intake Week 4 | Stress: (F(1,42) =0.1, p>0.05) | N/A |
| | | Drug: (F(1,42) =0.3, p>0.05) | N/A |
| | | Stress*Drug: (F(1,42) =0.1, p>0.05) | N/A |
| | **Water Intake Week 5** | Stress: (F(1,42) =0.1, p>0.05) | N/A |
| | | Drug: (F(1,42) =0.7, p>0.05) | N/A |
| | | Stress*Drug: (F(1,42) =1.5, p>0.05) | N/A |
| | Latency to Drink in Homecage | Stress: (F(1,42) =0.3, p>0.05) | N/A |
| | | Drug: (F(1,42) =0.2, p>0.05) | N/A |
| | | Stress*Drug: (F(1,42) =2.7, p>0.05) | N/A |
| | Latency to Eat in Homecage | Stress: (F(1,42) =0.1, p>0.05) | N/A |
| | | Drug: (F(1,42) =0.1, p>0.05) | N/A |
| | | Stress*Drug: (F(1,42) =3.3, p>0.05) | N/A |
| | Locomotor | Stress: (F(1,42) =1.9, p>0.05) | N/A |
| | | Drug: (F(1,42) =0.5, p>0.05) | N/A |
| | | Stress*Drug: (F(1,42) =1.0, p>0.05) | N/A |

**Table 3: Statistical analysis of the sex effects for the PFC GFAP+ cell activity enhancement study**

| Week of Testing | Behavioural Test | Statistics (F,p) | Post-hoc analysis |
|---|---|---|---|
| Baseline | Sucrose Consumption Test | Sex: (F(1,40) =2.6, p>0.05) | N/A |
| | | Sex*Stress: (F(1,40) =0.9, p>0.05) | N/A |
| | Water Test | Sex: (F(1,40) =0.1, p>0.05) | N/A |
| | | Sex*Stress: (F(1,40) =1.4, p>0.05) | N/A |
| | Coat State | Sex: (F(1,40) =5.8, p<0.05) | M>F* |
| | | Sex*Stress: (F(1,40) =1.3, p>0.05) | N/A |
| | Residual Avoidance | Sex: (F(1,40) =1.5, p>0.05) | N/A |
| | | Sex*Stress: (F(1,40) =1.5, p>0.05) | N/A |
| | Phenotyper - Time in Shelter | Sex: (F(1,40) =4.4, p<0.05) | M>F* |
| | | Sex*Stress: (F(1,12) =0.4, p>0.05) | N/A |
| | | Sex*Time: (F(1,12) =3.5, p<0.0001) | No difference between comparable groups were found |
| | | Sex*Time*Stress: (F(1,12) =1.5, p>0.05) | N/A |
| Week 1 | Sucrose Consumption Test | Sex: (F(1,40) =0.2, p>0.05) | N/A |
| | | Sex*Stress: (F(1,40) =0.1, p>0.05) | N/A |
| | Water Test | Sex: (F(1,40) =0.5 , p>0.05) | N/A |
| | | Sex*Stress: (F(1,40) =0.1, p<0.05) | N/A |
| | Coat State | Sex: (F(1,40) =35.7, p<0.0001) | See Sex*Stress |
| | | Sex*Stress: (F(1,40) =11.8, p<0.01) | Stress (M>F****), No Stress+M < Stress+M**** |
| | Residual Avoidance | Sex: (F(1,40) =1.6, p>0.05) | N/A |
| | | Sex*Stress: (F(1,40) =1.6, p>0.05) | N/A |
| | Phenotyper - Time in Shelter | Sex: (F(1,40) =4.0, p>0.05) | N/A |
| | | Sex*Stress: (F(1,12) =0.1, p>0.05) | N/A |
| | | Sex*Time: (F(1,12) =2.1, p<0.05) | 2 am - Stress (M>F*): 4 am - No Stress (M>F**) |
| | | Sex*Time*Stress: (F(1,12) =1.7, p>0.05) | N/A |
| Week 2 | Sucrose Consumption Test | Sex: (F(1,40) =0.1, p>0.05) | N/A |
| | | Sex*Stress: (F(1,40) =5.1, p<0.05) | No Stress+F >Stress+F****;No Stress+M >Stress+M** |
| | Water Test | Sex: (F(1,40) =0.3, p>0.05) | N/A |
| | | Sex*Stress: (F(1,40) =0.1, p<0.05) | N/A |
| | Coat State | Sex: (F(1,40) =17.6, p<0.0001) | M>F**** |
| | | Sex*Stress: (F(1,40) =2.0, p>0.05) | N/A |
| | Residual Avoidance | Sex: (F(1,40) =6.0, p<0.05) | F>M* |
| | | Sex*Stress: (F(1,40) =6.0, p<0.05) | No Stress+F >Stress+F****;No Stress+M >Stress+M**** |
| | Phenotyper - Time in Shelter | Sex: (F(1,40) =1.8, p>0.05) | N/A |
| | | Sex*Stress: (F(1,12) =0.4, p>0.05) | N/A |
| | | Sex*Time: (F(1,12) =0.6, p>0.05) | N/A |
| | | Sex*Time*Stress: (F(1,12) =1.3, p>0.05) | N/A |
| Week 3 | Sucrose Consumption Test | Sex: (F(1,36) =4.2, p<0.05) | F>M* |
| | | Sex*Stress: (F(1,36) =2.2, p>0.05) | N/A |
| | | Sex*Drug: (F(1,36) =0.1, p>0.05) | N/A |
| | | Sex*Stress*Drug: (F(1,36) =1.5, p>0.05) | N/A |
| | Water Test | Sex: (F(1,36) =1.3, p<0.05) | N/A |
| | | Sex*Stress: (F(1,36) =2.8, p>0.05) | N/A |
| | | Sex*Drug: (F(1,36) =0.1, p>0.05) | N/A |
| | | Sex*Stress*Drug: (F(1,36) =0.4, p>0.05) | N/A |
| | Coat State | Sex: (F(1,36) =47.7, p<0.0001) | M>F**** |
| | | Sex*Stress: (F(1,36) =7.6, p<0.01) | No Stress+No CNO: (M>F*); Stress+CNO: (M>F****); Stress+No CNO: (M>F****); No Stress+CNO+M < Stress+CNO+M***; No Stress+No CNO+F < Stress+NoCNO+F; No Stress+NoCNO+M < Stress+NoCNO+M**** |
| | | Sex*Drug: (F(1,36) =0.7, p>0.05) | N/A |
| | | Sex*Stress*Drug: (F(1,36) =0.4, p>0.05) | N/A |
| | Residual Avoidance | Sex: (F(1,36) =44.2, p<0.0001) | F>M**** |
| | | Sex*Stress: (F(1,36) =32.41, p<0.0001) | Stress+CNO: (F>M****); Stress+No CNO: (F>M****); No Stress+CNO+F < Stress+CNO+F****; No Stress+CNO+F < Stress+CNO+F****; No Stress+CNO+M < Stress+CNO+M* |
| | | Sex*Drug: (F(1,36) =0.1, p>0.05) | N/A |
| | | Sex*Stress*Drug: (F(1,36) =0.9, p>0.05) | N/A |
| | Phenotyper - Time in Shelter | Sex: (F(1,36) =2.1, p>0.05) | N/A |
| | | Sex*Stress: (F(1,36) =0.9, p>0.05) | N/A |
| | | Sex*Drug: (F(1,36) =0.2, p>0.05) | N/A |
| | | Sex*Stress*Drug: (F(1,36) =0.6, p>0.05) | N/A |
| | | Sex*Time: (F(1,12) =2.1, p<0.05) | See Sex*Time*Stress |
| | | Sex*Time*Stress: (F(1,12) =1.9, p<0.05) | 1am - Stress+No CNO: (F>M*); 3am - No Stress+No CNO: (M>F*), Stress+CNO: (M>F*); 4am - No Stress+No CNO: (M>F*), 5am- No Stress+No CNO: (M>F**) |
| | | Sex*Time*Group: (F(1,12) =1.4, p>0.05) | N/A |
| | | Sex*Stress*Drug*Time: (F(1,12) =1.3, p>0.05) | N/A |

| | | | |
|---|---|---|---|
| Week 4 | Sucrose Consumption Test | Sex: (F(1,36) =4.7, p<0.05) | M>F* |
| | | Sex*Stress: (F(1,36) =0.1, p>0.05) | N/A |
| | | Sex*Drug: (F(1,36) =0.2, p>0.05) | N/A |
| | | Sex*Stress*Drug: (F(1,36) =1.2, p>0.05) | N/A |
| | Water Test | Sex: (F(1,36) =1.2, p<0.05) | N/A |
| | | Sex*Stress: (F(1,36) =0.5, p>0.05) | N/A |
| | | Sex*Drug: (F(1,36) =0.1, p>0.05) | N/A |
| | | Sex*Stress*Drug: (F(1,36) =1.3, p>0.05) | N/A |
| | Coat State | Sex: (F(1,36) =22.1, p<0.0001) | M>F**** |
| | | Sex*Stress: (F(1,36) =3.9, p>0.05) | N/A |
| | | Sex*Drug: (F(1,36) =0.1, p>0.05) | N/A |
| | | Sex*Stress*Drug: (F(1,36) =0.1, p>0.05) | N/A |
| | Residual Avoidance | Sex: (F(1,36) =13.5, p<0.001) | F>M*** |
| | | Sex*Stress: (F(1,36) =18.6, p<0.0001) | Stress+CNO: (F>M****); Stress+No CNO: (F>M**); No Stress+CNO+F < Stress+CNO+F****; No Stress+CNO+F < Stress+CNO+F****; No Stress+CNO+M < Stress+CNO+M***; No Stress+CNO+M < Stress+CNO+M**** |
| | | Sex*Drug: (F(1,36) =0.1, p>0.05) | N/A |
| | | Sex*Stress*Drug: (F(1,36) =0.8, p>0.05) | N/A |
| | Phenotyper - Time in Shelter | Sex: (F(1,36) =3.4, p>0.05) | N/A |
| | | Sex*Stress: (F(1,36) =0.2, p>0.05) | N/A |
| | | Sex*Drug: (F(1,36) =0.1, p>0.05) | N/A |
| | | Sex*Stress*Drug: (F(1,36) =0.6, p>0.05) | N/A |
| | | Sex*Time: (F(1,12) =1.4, p>0.05) | N/A |
| | | Sex*Time*Stress: (F(1,12) =1.2, p>0.05) | N/A |
| | | Sex*Time*Group: (F(1,12) =1.9, p<0.05) | 7pm - Stress+No CNO: (F>M*); 3am - NoStress+CNO: (M>F**), NoStress+CNO+M > NoStress+NoCNO+M |
| | | Sex*Stress*Drug*Time: (F(1,12) =1.6, p>0.05) | N/A |
| Week 5 | Sucrose Consumption Test | Sex: (F(1,36) =1.2, p>0.05) | N/A |
| | | Sex*Stress: (F(1,36) =0.3, p>0.05) | N/A |
| | | Sex*Drug: (F(1,36) =0.2, p>0.05) | N/A |
| | | Sex*Stress*Drug: (F(1,36) =1.7, p>0.05) | N/A |
| | Water Test | Sex: (F(1,36) =0.4, p>0.05) | N/A |
| | | Sex*Stress: (F(1,36) =1.3, p>0.05) | N/A |
| | | Sex*Drug: (F(1,36) =1.8, p>0.05) | N/A |
| | | Sex*Stress*Drug: (F(1,36) =2.3, p>0.05) | N/A |
| | Coat State | Sex: (F(1,36) =33.9, p<0.0001) | M>F**** |
| | | Sex*Stress: (F(1,36) =2.1, p>0.05) | N/A |
| | | Sex*Drug: (F(1,36) =0.7, p>0.05) | N/A |
| | | Sex*Stress*Drug: (F(1,36) =0.1, p>0.05) | N/A |
| | Residual Avoidance | Sex: (F(1,36) =47.8, p<0.0001) | F>M**** |
| | | Sex*Stress: (F(1,36) =31.1, p<0.0001) | Stress+CNO (F>M**); Stress+No CNO (F>M**); No Stress+CNO+F < Stress+CNO+F****; No Stress+CNO+M < Stress+CNO+M**; No Stress+No CNO+F < Stress+ No CNO+F***; No Stress+No CNO+M < Stress+No CNO+M* |
| | | Sex*Drug: (F(1,36) =0.1, p>0.05) | N/A |
| | | Sex*Stress*Drug: (F(1,36) =1.4, p>0.05) | N/A |
| | Phenotyper - Time in Shelter | Sex: (F(1,36) =1.0, p>0.05) | N/A |
| | | Sex*Stress: (F(1,36) =1.8, p>0.05) | N/A |
| | | Sex*Drug: (F(1,36) =1.0, p>0.05) | N/A |
| | | Sex*Stress*Drug: (F(1,36) =1.8, p>0.05) | N/A |
| | | Sex*Time: (F(1,12) =1.9, p<0.05) | See Sex*Time*Stress |
| | | Sex*Time*Stress: (F(1,12) =2.6, p<0.01) | 10pm - No Stress+No CNO (M>F*); No Stress+No CNO+M < Stress+No CNO+M*, 5am - No Stress+CNO (M>F**); No Stress+No CNO(M>F*); No Stress+CNO+F < Stress+CNO+F*, No Stress+No CNO+F < Stress+No CNO+F* |
| | | Sex*Time*Group: (F(1,12) =0.7, p>0.05) | N/A |
| | | Sex*Stress*Drug*Time: (F(1,12) =0.5, p>0.05) | N/A |
| Week 6 | NIH | Sex: (F(1,36) =1.9, p<0.05) | N/A |
| | | Sex*Stress: (F(1,36) =0.4, p<0.05) | N/A |
| | | Sex*Drug: (F(1,36) =2.4, p>0.05) | N/A |
| | | Sex*Stress*Drug: (F(1,36) =2.2, p>0.05) | N/A |
| | NSF | Sex: (F(1,36) =0.9, p>0.05) | N/A |
| | | Sex*Stress: (F(1,36) =0.2, p>0.05) | N/A |
| | | Sex*Drug: (F(1,36) =0.4, p>0.05) | N/A |
| | | Sex*Stress*Drug: (F(1,36) =0.1, p>0.05) | N/A |
| | Locomotor | Sex: (F(1,36) =11.0, p<0.01) | F>M** |
| | | Sex*Stress: (F(1,36) =0.9, p>0.05) | N/A |
| | | Sex*Drug: (F(1,36) =0.1, p>0.05) | N/A |
| | | Sex*Stress*Drug: (F(1,36) =0.7, p>0.05) | N/A |
